# MAJORA: Continuous integration supporting decentralised sequencing for SARS-CoV-2 genomic surveillance

**DOI:** 10.1101/2020.10.06.328328

**Authors:** Samuel M. Nicholls, Radoslaw Poplawski, Matthew J. Bull, Anthony Underwood, Michael Chapman, Khalil Abu-Dahab, Ben Taylor, Ben Jackson, Sara Rey, Roberto Amato, Rich Livett, Sónia Gonçalves, Ewan M. Harrison, Sharon J. Peacock, David M Aanensen, Andrew Rambaut, Thomas R. Connor, Nicholas J. Loman, The COVID-19 Genomics UK (COG-UK) Consortium

## Abstract

Genomic epidemiology has become an increasingly common tool for epidemic response. Recent technological advances have made it possible to sequence genomes rapidly enough to inform outbreak response, and cheaply enough to justify dense sampling of even large epidemics. With increased availability of sequencing it is possible for agile networks of sequencing facilities to collaborate on the sequencing and analysis of epidemic genomic data.

In response to the ongoing SARS-CoV-2 pandemic in the United Kingdom, the COVID-19 Genomics UK (COG-UK) consortium was formed with the aim of rapidly sequencing SARS-CoV-2 genomes as part of a national-scale genomic surveillance strategy. The network consists of universities, academic institutes, regional sequencing centres and the four UK Public Health Agencies.

We describe the development and deployment of Majora, an encompassing digital infrastructure to address the challenge of collecting and integrating both genomic sequencing data and sample-associated metadata produced across the COG-UK network. The system was designed and implemented pragmatically to stand up capacity rapidly in a pandemic caused by a novel virus. This approach has underpinned the success of COG-UK, which has rapidly become the leading contributor of SARS-CoV-2 genomes to international databases and has generated over 60,000 sequences to date.

## Introduction

Combining genomic sequencing of pathogens with epidemiology as part of a response to an outbreak has demonstrated success in epidemiological investigations of viruses such as Ebola, Yellow Fever and Zika [1]. Pathogen genomes are useful for reconstructing a phylogenetic history of an outbreak, and are now being used in real-time to assist epidemic response.

Established sequencing networks already exist for some infectious pathogens. As an example, the GenomeTrakr Network is part of the US Food and Drug Administration and connects labs across the United States and internationally to sequence foodborne bacterial pathogens and since 2013 the project has sequenced nearly 500,000 isolates. Flu viruses are also routinely sequenced, both through the use of Sanger and NGS techniques. In the UK, Public Health England and Public Health Wales both operate seasonal influenza surveillance programmes using NGS, with results reported to both Governments and international organisations such as the WHO and ECDC. Increasingly, public health agencies are developing surveillance programmes for viruses that are built on the use of NGS data. While genomic data is increasingly used within public health agencies for retrospective surveillance activities, the benefits of genomic epidemiology are yet to be fully realised for prospective and proactive outbreak response. This is exemplified in the current pandemic, where the initiation of programmes for sequencing of SARS-CoV-2 typically lagged behind planning for other parts of the pandemic response. The utility of genomic data has been such that this should be the last pandemic where genomic epidemiology is not a core part of pandemic planning.

Most existing public health sequencing initiatives are built around whole genome sequencing capacity afforded by facilities in large hospitals and public laboratories. However, with the emergence of lower cost sequencing instruments such as Oxford Nanopore, genomic sequencing is now available to smaller regional hospitals and academic laboratories, vastly expanding the sequencing capacity for a hypothetical surveillance network. Such technology is small and cost-effective enough to conduct sequencing of small pathogen genomes in the field, in the clinic and in the classroom. However, with this democratisation of sequencing technologies, a new challenge emerges in how data generated across many different laboratories can be collated, compared and analysed to support outbreak/pandemic response simultaneously at local, regional, national and global levels.

The COVID-19 Genomics UK (COG-UK) consortium was established in March 2020 with the aim to deliver large-scale and rapid whole-genome virus sequencing and analyze the sequences for local NHS centres and the UK government [2]. COG-UK is a national partnership of NHS organisations, the four UK Public Health Agencies, the Wellcome Sanger Institute and over 20 academic partners. The work of the consortium generates reports for the UK Scientific Advisory Group for Emergencies (SAGE), as well as providing analyses and advice to the UK devolved administrations, such as via the Welsh Government Technical Advisory Group. This is the first time that genomic epidemiology has been used at a national scale to guide a response to a pandemic in the United Kingdom, as demonstrated in regular reports to the UK’s Scientific Advisory Group for Epidemics (https://cogconsortium.uk/news).

As well as rapidly responding to the problems of how to extract and sequence SARS-CoV-2 genomes, another key challenge for COG-UK was how to develop an infrastructure capable of harmonising data from a network of sources to create one dataset for analysis. The development of this system posed many interesting and challenging problems from a technical standpoint. In this article we present several such problems, our solutions and what we have learned from the process. Our system provides a model (Figure 1) that may serve as a foundation to inform others who are faced with the challenge of designing and deploying a similar system to aid outbreak tracking in this or future pandemics.

**Figure 1.**
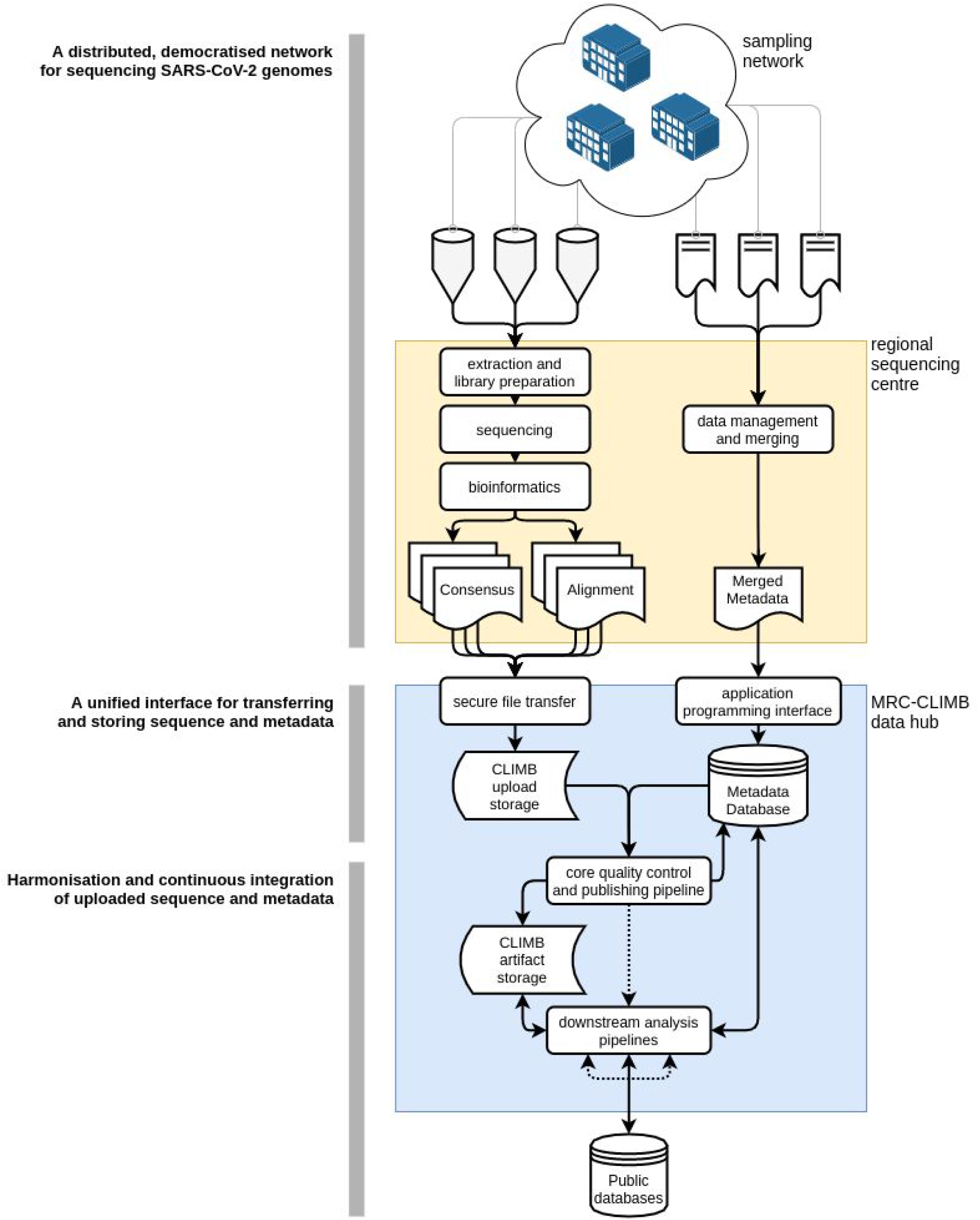
Overview of the COG-UK data flow. (Top) A network of sampling sites (hospitals, GPs, etc.) produce samples and sample metadata which are received by a regional sequencing centre. The sample is extracted and sequenced and a locally run bioinformatics pipeline generates both a consensus viral genome and an alignment of sequenced read fragments against the SARS-CoV-2 reference genome. (Middle) The consensus sequence and alignments are uploaded via secure file transfer to be stored on MRC-CLIMB. Metadata is securely transferred over HTTPS to an application programming interface (API) that transforms metadata into a model to be stored in a database on MRC-CLIMB. (Bottom) The core quality control pipeline executes every day to integrate newly uploaded samples and metadata into the single canonical dataset of all uploaded sequences. Once this pipeline is finished, it notifies downstream analysis pipelines through a messaging protocol to generate analysis artifacts like phylogenetic trees. Downstream analysis pipelines also automatically deposit genomes in public databases such as GISAID and INSDC/ENA.

## Results

We present a model of our system (Figure 1), which can be broken down into three core functions:

- Produce data, by connecting a network of regional sequencing sites (academic or government affiliated) to a network of sampling organisations, to establish a distributed, democratised network for sequencing SARS-CoV-2 genomes
- Collect data by providing a system to transfer sequencing data, consensus genomes and sample metadata that works in the same way for every member of the consortium
- Integrate data into a single dataset by harmonising the collected sequences and metadata

### An autonomous and scalable network for decentralised sequencing of SARS-CoV-2 genomes

The COG-UK consortium forms a national network of organisations that in combination collect and sequence samples. The organisations within the consortium have a high degree of autonomy. This autonomy is valuable as sites can take advantage of their own local expertise to make decisions on protocols and methods to use for sample collection, preparation and sequencing; reducing the burden for an organisation that wishes to participate. Some of these sampling sites also have the capacity and resources to also perform their own sequencing, those that do not are connected to either a regional sequencing organisation, or the Wellcome Sanger Institute (WSI). Regional sequencing sites include academic institutions, small laboratories and public health agencies. Connecting sampling organisations to a local sequencing laboratory means sequenced genomes can be turned around within 24-48 hours of sample collection.

This two-tiered sequencing model has facilitated both a prioritised, rapid regional response; as well as supporting lower priority, high-throughput projects such as the sequencing of every positive sample from the Lighthouse Laboratories.

However, this autonomy comes at a cost: raising the difficult challenge of coordinating such a diverse network of sites; using a spectrum of methods for sample extraction, PCR, library preparation, sequencing and consensus-generating bioinformatics. The core problem we faced when tasked to build this infrastructure is ultimately one of data interoperability. With so many geographically dispersed sequencing operations and the four public agencies all producing data with a wide variety of different techniques and platforms, it was necessary to deploy an infrastructure to collate this data into a single, consistent, canonical data set, available for everyone within the consortium.

#### A hub model for integrating genomic and epidemiological data

We chose to form a hub model around the Cloud Infrastructure for Microbial Bioinformatics (CLIMB) compute facility [3]. CLIMB is not just a pragmatic choice given the affiliation of the authors, since first deployed in 2014, it has provided infrastructure to microbiologists to produce and use software for the analysis of genomic data sets, serving over 300 research groups at more than 85 organisations spread across the United Kingdom (including PHE and PHW). It was designed as a system to support microbial bioinformatics and has been used for pathogen outbreak analysis in the past [4].

We isolated over 600 vCPUs of compute from the CLIMB infrastructure to form a ‘walled garden’ system named CLIMB-COVID, which was deployed in three days for the purpose of providing a central, replicated environment for the storage and analysis of data generated by COG-UK. Additionally, CLIMB offered a third party neutral territory; as it is external to any specific university, government or public health agency governance. This in turn has enabled cooperation across a diverse network of sequencing operations, and the development of a bespoke service and environment to meet the needs of the project.

Sites participating in the consortium maintain authority over the data they generate, interpreting and sharing it to inform a local public health response. As part of their membership they are responsible for transferring the sequenced consensus FASTA file, and an alignment of the sequenced reads against the SARS-CoV-2 reference genome [5] as a BAM to a designated server hosted on the CLIMB.

To assist with the on-boarding of new sites, including those with limited bioinformatics support we also built a reproducible Nextflow pipeline (https://github.com/connor-lab/ncov2019-artic-nf) that enables the processing of data for sites following the ARTIC protocols.

#### A walled garden for fast turn around and to maintain sequence integrity

This hub model operates with a different paradigm to one suggested recently by Black et al. [6], which recommended that raw reads would first be uploaded to the SRA, or Illumina BaseSpace; and that the final step of any assembly pipeline would be automatic submission to a public database such as one of the International Nucleotide Sequence Database Collaboration (INSDC) databases, or a pathogen specific initiative such as GISAID. However this paradigm would introduce unnecessary delays in the processing of data and hamper real-time genomic surveillance efforts. In building our system the focus has been on generating actionable information to support public health action as rapidly as possible.

Our approach instead takes sequence data for initial analysis inside a system hosted on MRC-CLIMB which can only be accessed by members of the consortium. This ensures the data is immediately usable, as sequences can be transferred to the consortium as soon as they have been processed locally; whereas large public databases often have a lead time up to a few days before accessions are indexed and resources can be downloaded, which is incompatible with the goal to turn around sequences within 24 hours.

Having a walled garden system with defined entry points for data has enabled us to define the files to be uploaded, which includes consensus sequences and BAM files of read alignments as opposed to raw sequencing reads. This simplifies analysis within CLIMB-COVID, and also enables the hub to avoid storing human reads sequenced incidentally as part of SARS-CoV-2 sequencing, while also providing valuable data that can be used to perform additional analyses for scientific or quality control purposes.

This model also allows our internal pipelines to be tolerant of the different error profiles we may expect to see given the diverse methodologies in use across the sites. Uploading the data centrally allows us to perform basic quality control and ensure consensus genomes are internally consistent before they are distributed outside the consortium, mitigating the risk of polluting international databases. Requiring data to be uploaded to CLIMB-COVID centrally also enforces an environment that fosters data sharing, as sequences can only be analysed and integrated into the data set if they have been shared with the consortium.

#### Single, unique, shareable, perpetual identifiers for a centralised sample registry

A necessary component of the hub model is maintaining a registry of all sample identifiers sequenced by the consortium. To avoid confusion and aid interoperability, these identifiers should be unique and refer to one specific sample, from a particular sampling event. Identifiers should not encode any metadata that could be misinterpreted or would limit the ability to share the identifier with others. A sample should have a suitable identifier assigned as soon as practically possible, and it should never be changed or re-used in future.

These requirements are often easily violated in practice. Many samples in a diagnostic laboratory setting are automatically assigned a sequential number, or assigned a number which encodes information about the patient. This means that an identifier could contain personal identifiable information, may not be unique within the consortium, or would need additional metadata to be shared to allow for disambiguation. Even when PII is not present in the identifier, sample identifiers generated by public health agencies or hospitals are often used for linkage purposes, and may be considered privileged information, limiting their usability.

To overcome this, we used pre-printed barcodes, that could be distributed to collecting centres and affixed to a sample tube as soon as practically possible. They are easy to organise and distribute, cause little additional burden to laboratories, and ensures that each specimen receives an identifier that is safe to share with the consortium. The overhead of this solution is a linkage table is required to map the identifier of the specimen inside the collecting organisation, to the COG-UK identifier on our barcode label. This linkage holds the key for accessing more detailed information held about a sample by public health agencies at a later date. In many cases, the collecting organisation was confident that their identifiers were safe to share as part of our standard collection process. Where this is not the case, the linkage is held separately at the laboratory, creating a small hurdle for obtaining linkage.

Although the pre-printed barcodes simplify the process of generating identifiers for samples, it was more difficult to deploy a scheme for uniquely identifying the patients from which the samples were taken from. Governments and health agencies around the world have a single, unique, perpetual identifier for a citizen, but they are not shareable in practice as their inclusion makes the dataset patient identifiable information. We considered using a pseudo-identifier derived from these, e.g. based on a salted hash of the identifier, but the additional operational complexity was considered too great for local laboratories with limited bioinformatics capacity. In the scenario where a sample is known to have come from a patient who has already been sampled, we settled for using the identifier of the first sample as the patient identifier. This has limitations, but is readily achievable in cases where local laboratories know whether a sample has been collected from a patient with previous samples, and at least allows for grouping of samples from the same patient when that relationship is known.

#### A minimal metadata standard to ensure wide adoption of data collection

For the sequenced genomes to be useful, it is essential to pair them with metadata that contextualises the time, place and circumstance of the collected sample. This context is what allows us to use genomic epidemiology to drive an effective intervention as part of a public health response. Without adequate metadata, sequencing genomes is just an expensive form of stamp collecting.

There are already several well defined lists of metadata that are recommended for collection, for example submissions to the European Nucleotide Archive suggest following the “ENA virus pathogen reporting standard checklist” (ERC000033), and recently the Public Health Alliance for Genomic Epidemiology (PHA4GE) drafted a specification for sharing contextual data about SARS-CoV-2 genomes to advocate the openness and reusability of generated data sets [7].

Although it is straightforward to construct a list of desired pieces of metadata to collect, the real problem is reconciling such a standard with the reality of how data can be collected on the ground.

Until such a time where a standard methodology, legal framework and technical infrastructure for data sharing between various public health agencies, hospital trusts and academic laboratories exists, there is a need for pragmatism. We must expect scenarios where limited metadata will have been collected in the first place, or a diagnostic laboratory may not have the staff or resources to provide enough metadata to meet the requirements of a checklist. There will likely be a need for data sharing agreements to be drawn that will prohibit sharing some metadata fields more widely. Even in the case where a wealth of metadata is available, there may be a need to prioritize some fields over others; as multiple fields together may permit those fields to be used in conjunction with other data sources to aid identification of a person (deductive disclosure).

There is a balance to be struck to choose fields that can be practically collected by all organisations that join the consortium, and obtain enough metadata to do meaningful analysis. We defined a very small set of mandatory fields (Table 1) that aimed to limit the burden on laboratories and ensure that basic metadata was provided in a timely fashion. For a full table of fields refer to Table 3.

**Table 1.** COG-UK minimally useful metadata standard.

| **Data item** | **Field name** | **Description** | **Access level**<br>**(Public,**<br>**Consortium or**<br>**Restricted)** | **Mandatory** | **Implementation** |
| --- | --- | --- | --- | --- | --- |
| **Central Sample ID** | central_sample_id | A unique identifier to refer to the sample within the consortium | Public | Yes | Tier 1 database model (biosample) |
| **Date of sample (collected)** | collection_date | The date the sample was collected | Public | Yes (otherwise received_date) | Tier 1 database model (biosample collection) |
| **Date of sample (received)** | received_date | The earliest date that this sample was known to be checked in to a diagnostic or sequencing laboratory | Public | No (unless collection_date is not provided) | Tier 1 database model (biosample collection) |
| **UK nation** | adm1 | The country within the United Kingdom in which the sample was collected | Public | Yes | Tier 1 database model (biosample collection) |
| **County** | adm2 | The county within the United Kingdom in which the sample was collected | Consortium | Strongly recommended | Tier 1 database model (biosample collection) |
| **Sampling strategy** | is_surveillance | Whether this sample was collected as part of a random surveillance strategy, or a targeted outbreak analysis | Consortium | Yes | Tier 2 database model (COG-UK biosample collection supplement) |
| **Table 1 COG-UK minimally useful metadata standard** |  |  |  |  |  |

**Table 2.** Three tiers of metadata within Majora. Majora stores submitted metadata about artifacts and processes in an SQL database. Metadata is stored differently based on its priority. Fields that are a core part of a model (for example, a sample identifier, or the name of a file) are considered primary metadata and are stored in a distinct database model. Metrics such as the results of a PCR Ct test, or the coverage levels of a BAM are also stored in a distinct database model and are attached to primary models through a database foreign key. Arbitrary metadata can then be stored in key value pairs (not backed by any particular database model) and tagged to primary and secondary models as appropriate.

| Tier | Implementation | Properties | Example |
| --- | --- | --- | --- |
| Primary | Database model | Fast queries via object-relational mapping<br>Takes up space in database even if unused<br>Significant work to add to the database model, API and user templates | Biosample identifier<br>Patient sex, age<br>Digital resource file path, size, hash |
| Secondary | Database model | Fast queries via object-relational mapping<br>Additional lookups necessary to link back to the primary database model<br>Cannot assume a primary model will have a supplement | Cycle threshold metrics for biosamples<br>BAM coverage metrics<br>Patient healthcare worker or care home status |
| Tertiary | Key-value row in generic model | More difficult to manage artifacts based on tagged properties alone<br>Highly flexible<br>No work required to add new tags at any time | Locally relevant tags not implemented in a model<br>Additional anonymised patient information<br>Additional sequencing run information |

**Table 3.**
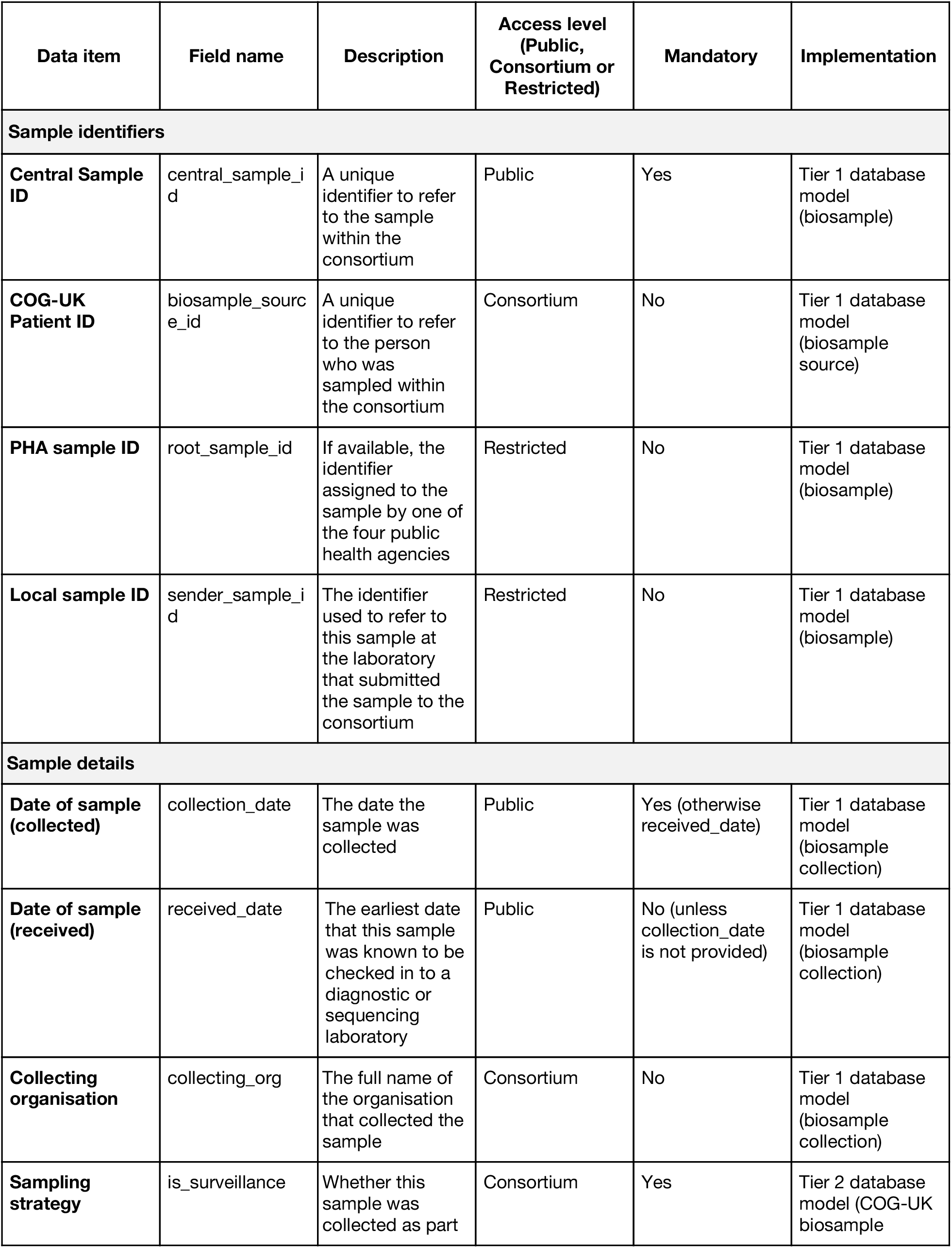
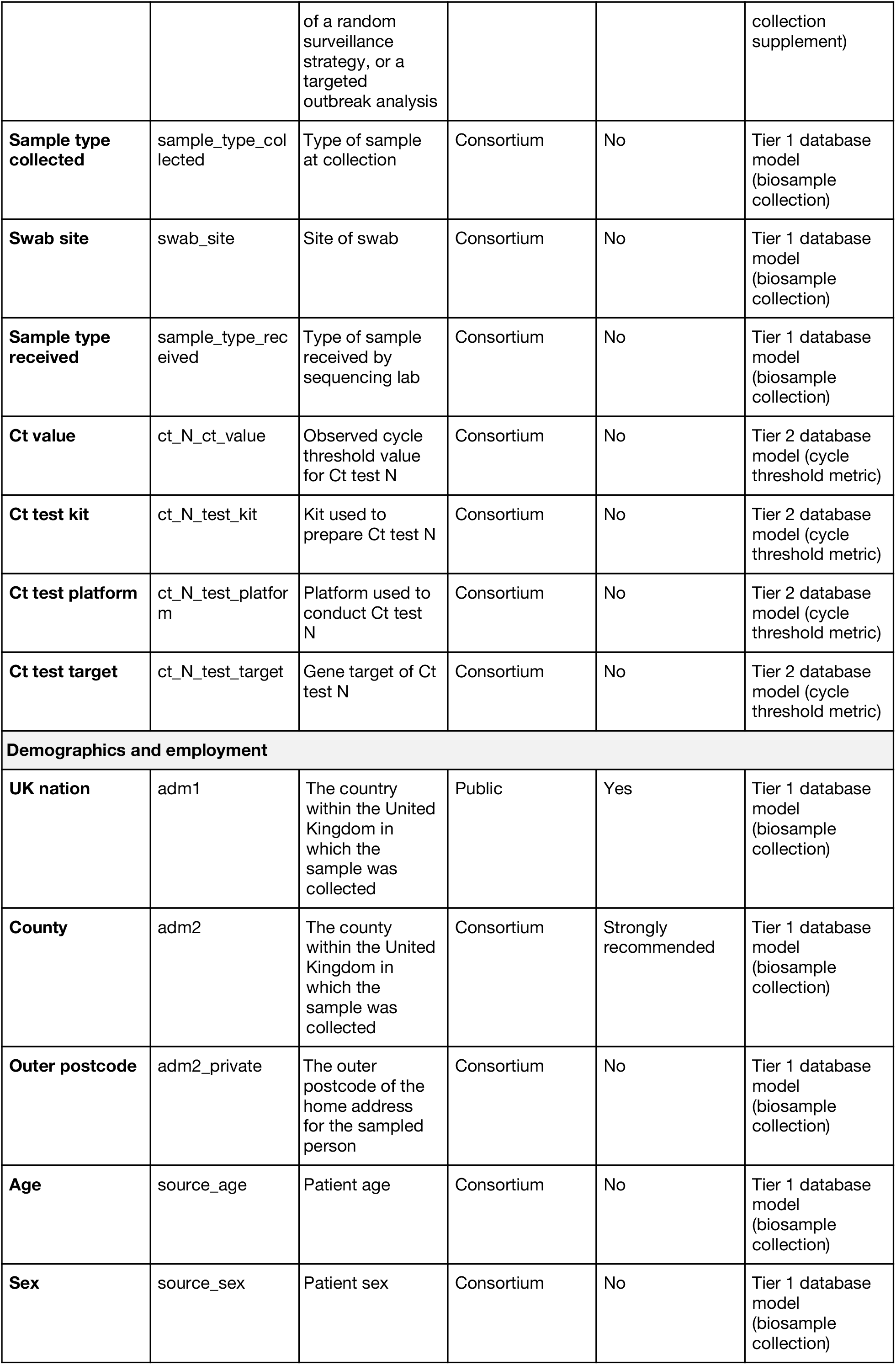
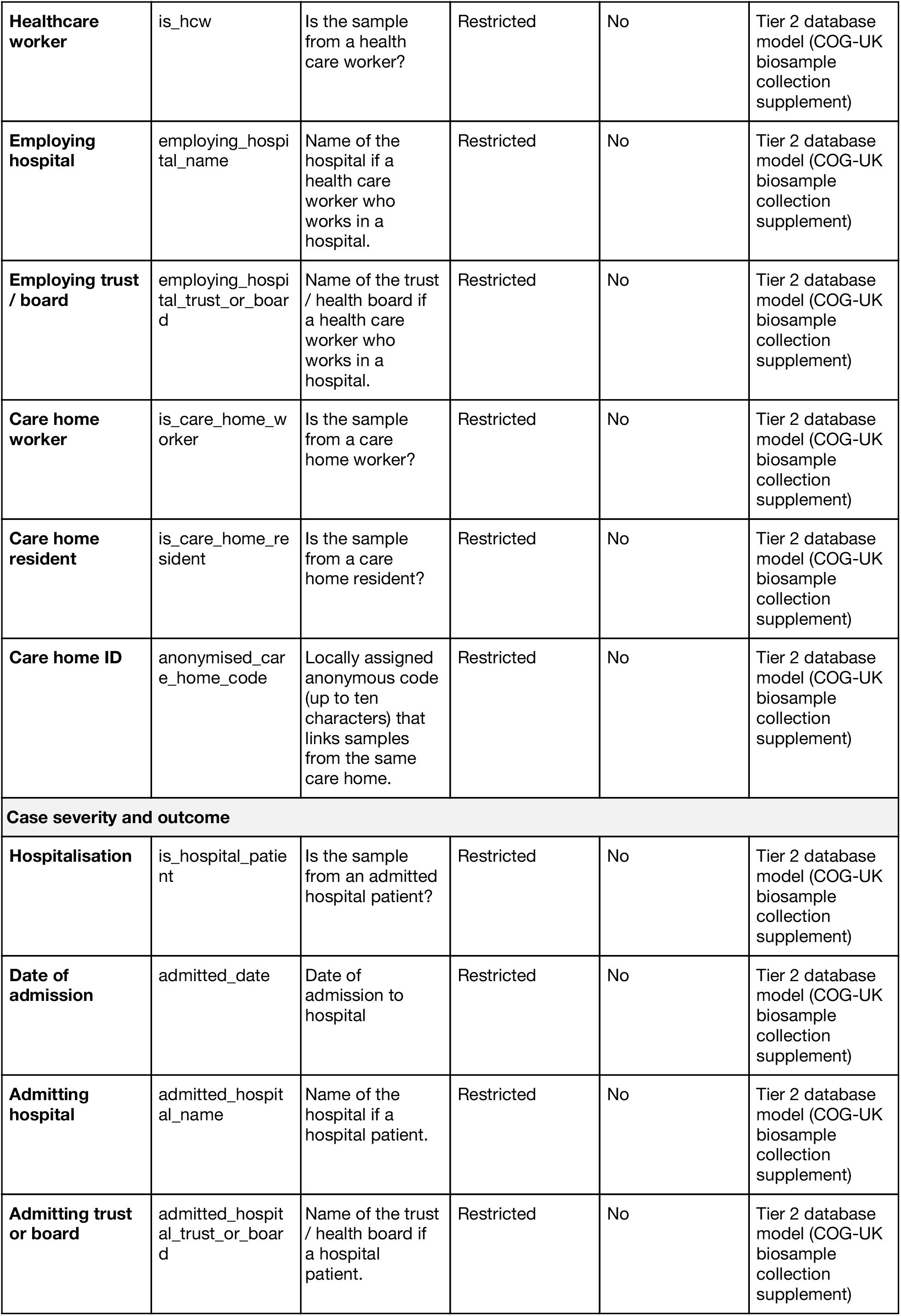
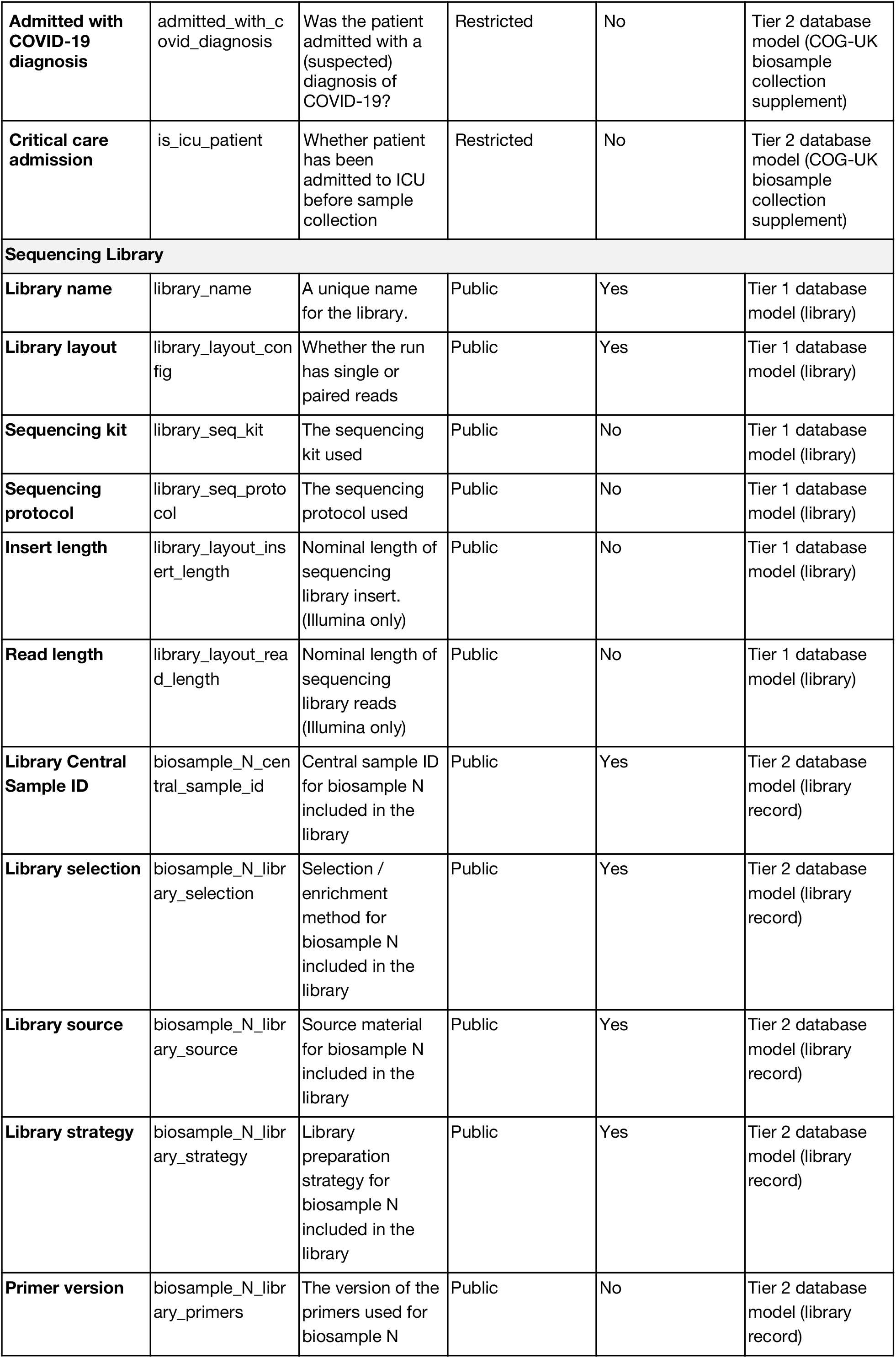
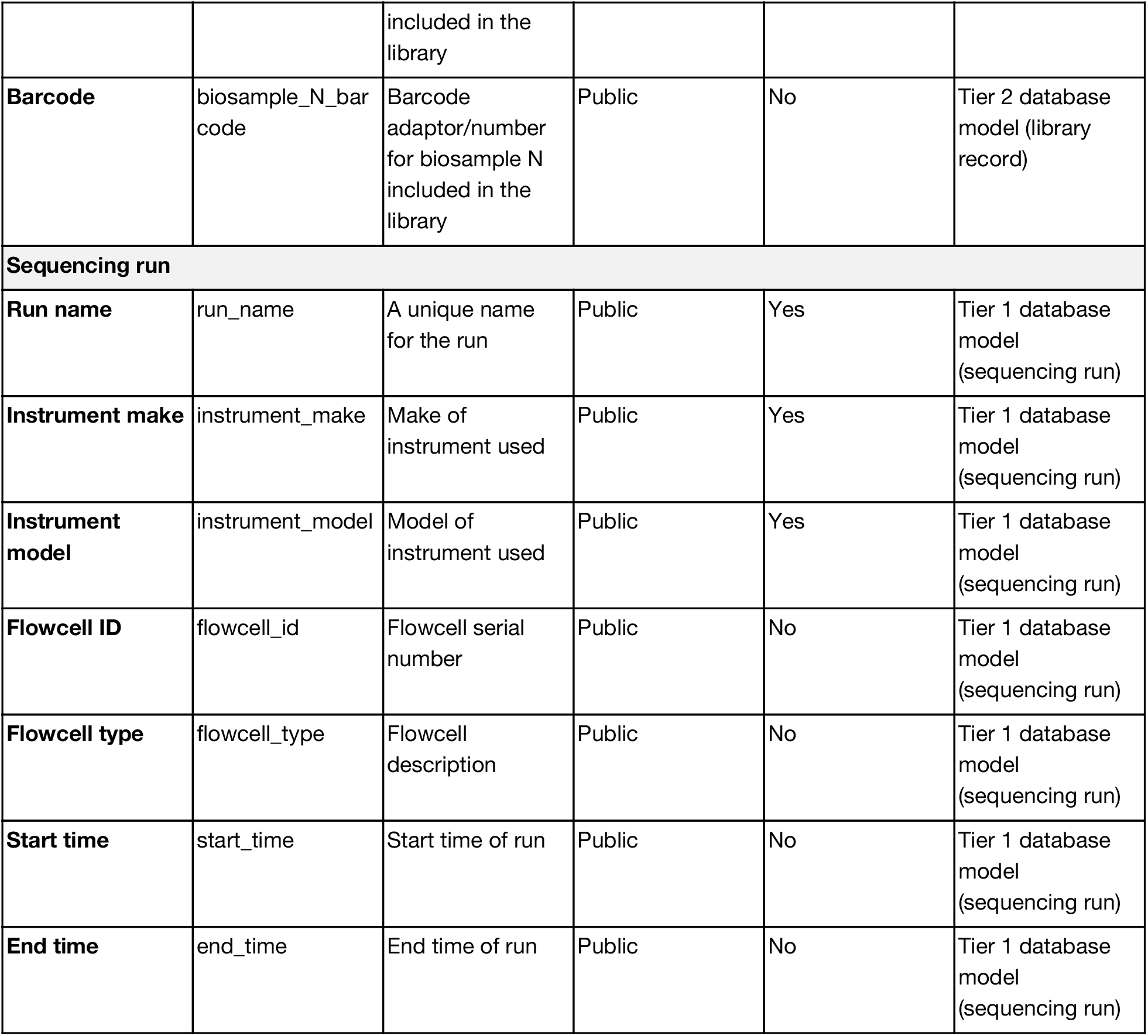
COG-UK full metadata standard.

The expertise of the analyst groups within COG-UK is focussed on viral phylodynamics, which looks to map how the evolution of sequenced viral genomes is linked to where and when those samples were collected. Our mandatory metadata fields reflect this, linking the date a sample was collected and the approximate geographical location it was collected in. County was a necessary but unfortunate compromise as the security assessments and contractual arrangements to collect more fine-scale location information such as outer postcode would take some time to organise.

Diagnostic laboratories are required to inform public health agencies when they identify notifiable organisms, including SARS-CoV-2. We have used this process to allow retrieval of additional data relating to the patient from whom the sample was taken. Laboratories either add the COG-UK identifier directly to the relevant national data systems or provide this separately to the public health agency. Again, a pragmatic approach has been taken with a variety of secure transfer methods used to provide this information as systems have matured. This has allowed the majority of samples to be linked to records held by public health agencies, who can provide supplementary metadata where required and use this information in their own analyses.

### A unified interface for transferring and storing sequences and sample metadata

#### A unified template for metadata collection

It was clear from the outset that we needed to provide a unified method for metadata to be uploaded, regardless of whether users are providing the minimal metadata or filling in every field, otherwise it would be impossible to maintain a centralised sample registry. Given the diverse nature of informatics skills across the participating organisations, a tool to upload metadata would need to be easily accessible (behind a variety of firewalls) and not require installation or configuration of software on the user’s machine.

At the middle of the intersection between intuitive and portable, spreadsheet software appears to have become a *de facto* method for clinicians and laboratory technicians to tabulate and share metadata. This is somewhat convenient, as spreadsheet software is almost always installed on computers used in diagnostic laboratories and can still generate highly portable comma or tab delimited versions of tabulated data. Leveraging this, we developed a CSV template of our minimal metadata standard. The template and associated documentation was iteratively developed in response to the needs of both the analysts and the feedback from participating sites on what can be realistically collected in a short amount of time. Organisations download a fresh copy of the template when samples are packaged and sent for sequencing and fill in the mandatory fields, and as many optional fields as they can.

Providing this intuitive way for users to provide data raises the issue of how these spreadsheets can be error checked, collated and made accessible to analysts. It is insufficient to hope that users will adhere to any validators built-in to the template, and some software may not support included validators. Once validated, one may consider just merging all these sheets together as a naive database. This may be reasonable in a small scale project, but for a robust national real-time response, this data needs to be supported by infrastructure capable of enforcing validation and adding new samples immediately.

#### Centrally managing consortium data through application programming interfaces (APIs) and Majora

It would not be feasible to manually moderate uploaded metadata, so we needed a system to alert users about invalid data and allow us to integrate valid metadata about samples into our dataset as soon as possible. There should be no human intervention involved in validating, processing, or querying metadata. To this end, a set of application programming interfaces (APIs) was developed. An API allows a computer program to interface with a human, or other computers. Within COG-UK, metadata can be submitted and queried through a collection of API endpoints exposed by a program named “Majora”, from which our whole system model gets its name.

The instance of Majora deployed for CLIMB-COVID was developed specifically for COG-UK, and is the brain of our digital infrastructure. It stores information about samples and files; which are referred to as “artifacts”. Majora also concerns itself with storing information on the “processes” that have been applied to these artifacts. For example, a group of sample artifacts may be pooled to form a library, a library is sequenced to provide signal data. Bioinformatics pipelines convert signal to reads, and reads to consensus genomes, and so on. By storing a record of how each artifact comes into being, and how artifacts are linked together through processes, it is possible to traverse a tree of processes, providing a full audit trail from when a sample was collected to any files and analyses generated about it downstream.

Majora (https://github.com/SamStudio8/majora/) is a Django application [8], with a set of APIs, a database of bespoke models and is also a web application in its own right. Users are able to register for an account that must be approved by their organisation’s lead user. The website allows for easy access to limited metadata and shows the history of processes that are known about a sample (Figure 4). For savvy users, bots and pipelines, a command line client has been developed that uses the API to access more advanced functionality.

**Figure 2.**
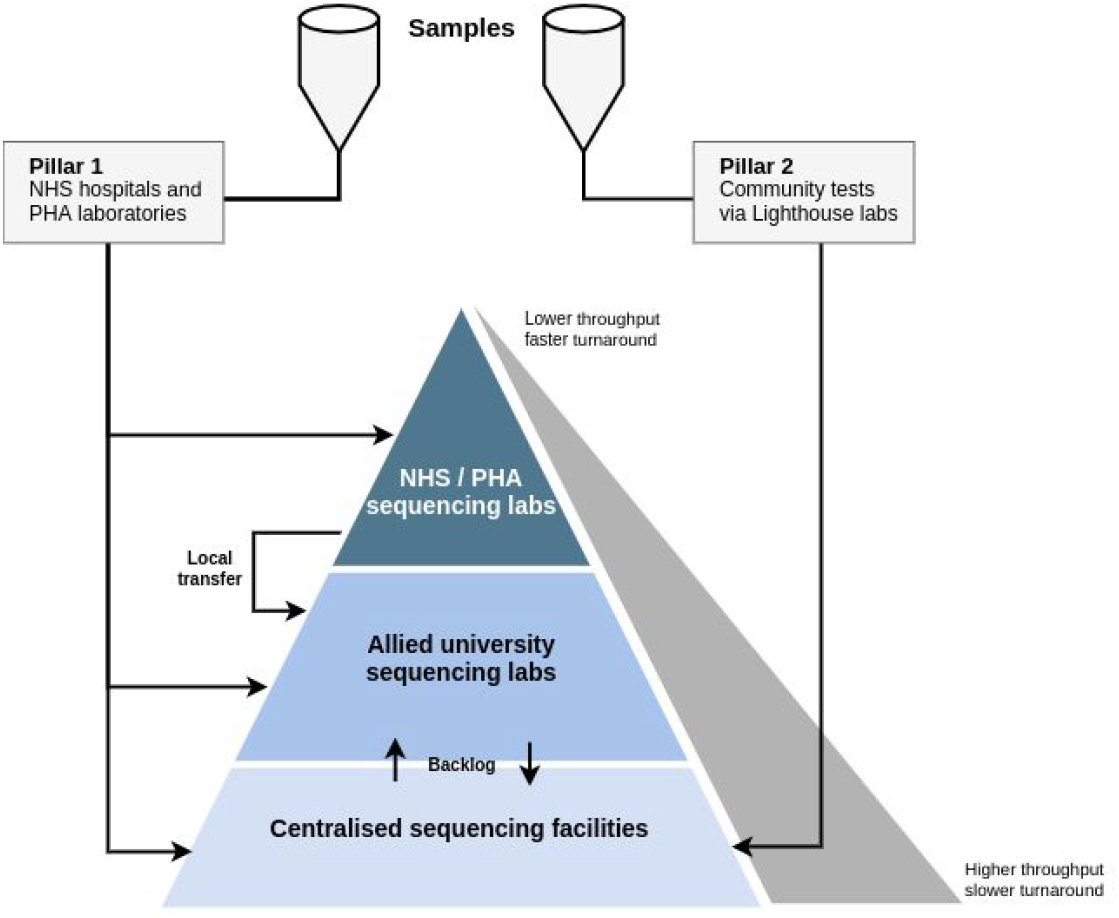
COG-UK sequencing model. Samples are sourced from two “pillars”; Pillar 1 samples are collected across the NHS and Public Health Agencies, Pillar 2 samples are collected at the Lighthouse Labs at particular strategic sites in the UK. Generally, Pillar 1 samples are received by NHS labs who process them for sequencing locally, or by a university sequencing lab for a fast turnaround. Pillar 2 samples are shipped through to the Wellcome Sanger Institute for high-capacity sequencing.

**Figure 3.**
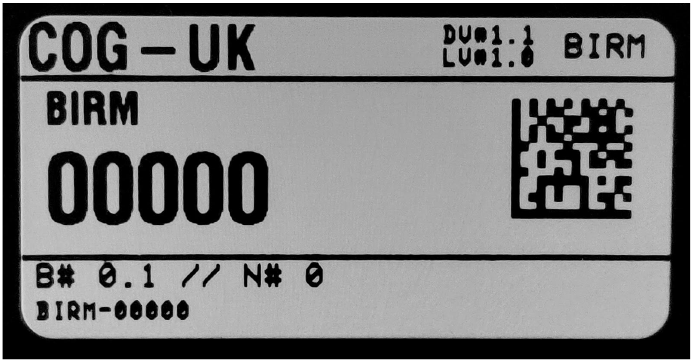
Example COG-UK sample label. A reference example of a label affixed to samples within the COG-UK consortium. The label is both human and machine readable. This is one of many label formats, owing to the decentralised nature of the consortium.

**Figure 4.**
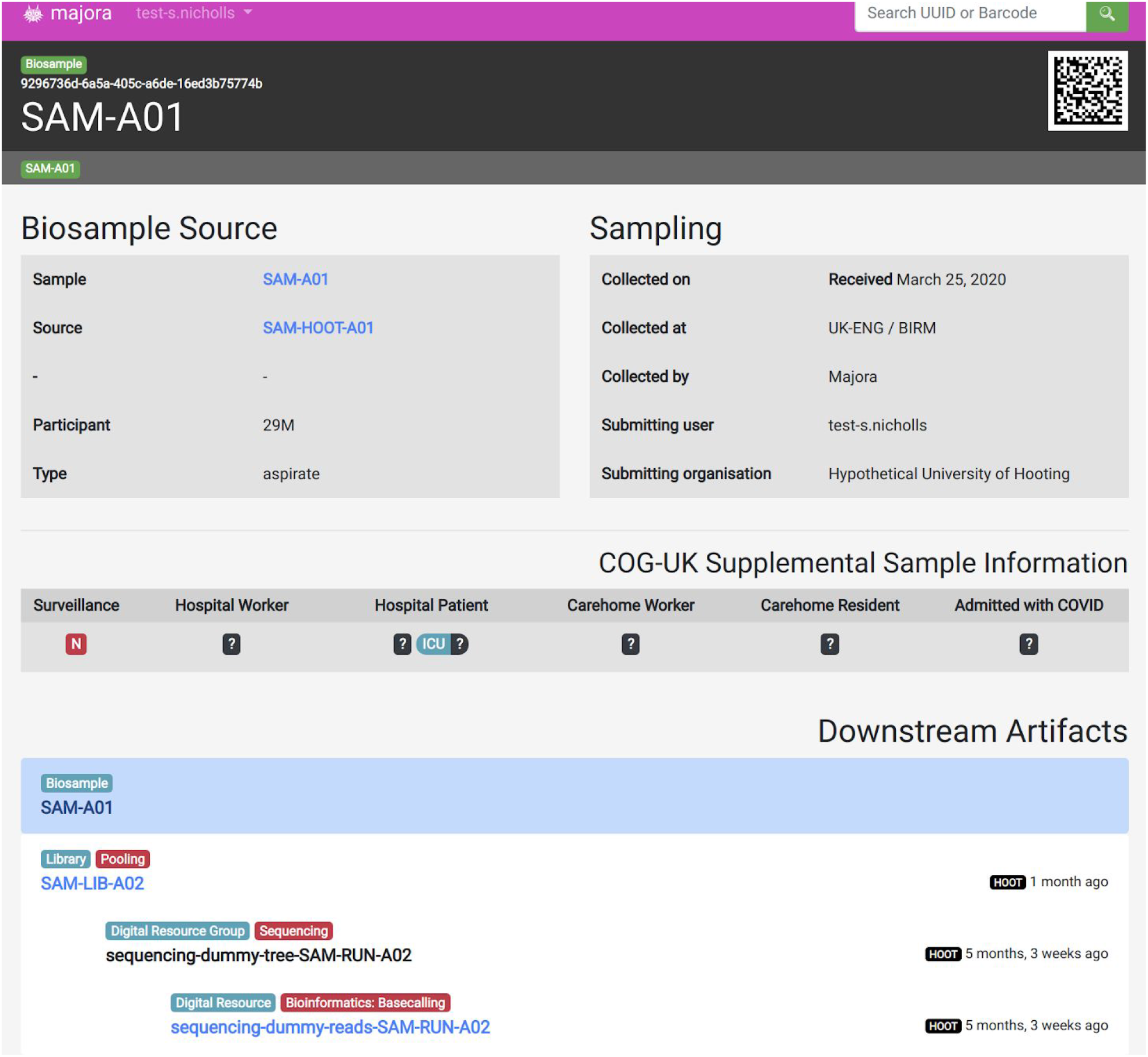
Majora web application biosample view. An example of the web interface presented by Majora to detail a biosample artifact. The downstream artifacts section allows users to see what processes have been applied to the biosample. In this example, the sample was incorporated onto a pooled sequencing library, which was sequenced and basecalled.

The website is protected by enforcing two factor authentication on users who wish to view any metadata, or use the APIs. The Majora APIs were initially secured with a rotating API key scheme that allowed external applications to perform actions as a user, without the user having to provide their actual account password. More recently, we are migrating away from these keys to a more straightforward, industry-standard protocol for authorization (OAuth 2.0).

As Majora is the only interface a user has to the metadata stored by the consortium, we have the opportunity to store restricted data and control access to it. As part of this control, we have integrated a system into Majora whereby data agreements can be viewed and signed by consortium users. For example, users are able to upload the “local” identifier of a sample as it is referred to inside of the collecting site (which is considered to be restricted), but we do not share this publicly, or within the consortium. Through Majora’s agreements system, users can give permission for the identifiers they have uploaded to be shared specifically with public health agencies, allowing COG-UK sequences to be linked to wider health informatics data. This layer between the users and the database where metadata is stored allows us to maintain an audit trail of who performed what actions both on the website, and through the API; satisfying the requirements set out by NHS digital.

The teams who determine the minimal metadata, define the templates and the developer of the API are closely linked, allowing the metadata collection strategy and the API to evolve with the changing demands of the consortium.

#### A user-friendly method for uploading and validating metadata

The COG-UK consortium includes hundreds of individuals working across tens of sites, with over 100 users registered to access CLIMB-COVID itself. Developing and deploying robust and well-documented APIs is therefore critical for reducing human workloads. Because of the size and scale of COG-UK it is critical to plan for scenarios where an API must be able to interact with a human in a friendly way, to ensure that the system can be supported by what remains a small team. For example, informing a user who has uploaded metadata that contains inconsistencies which need their attention. The APIs for uploading metadata require the fields to be arranged in a structured text format called JSON (JavaScript Object Notation) (Figure 5). Messages and validation errors are returned to the API user in the same format. Although JSON can be viewed in basic text readers, or pretty printed on a command line, they are not intended for human consumption.

**Figure 5.**
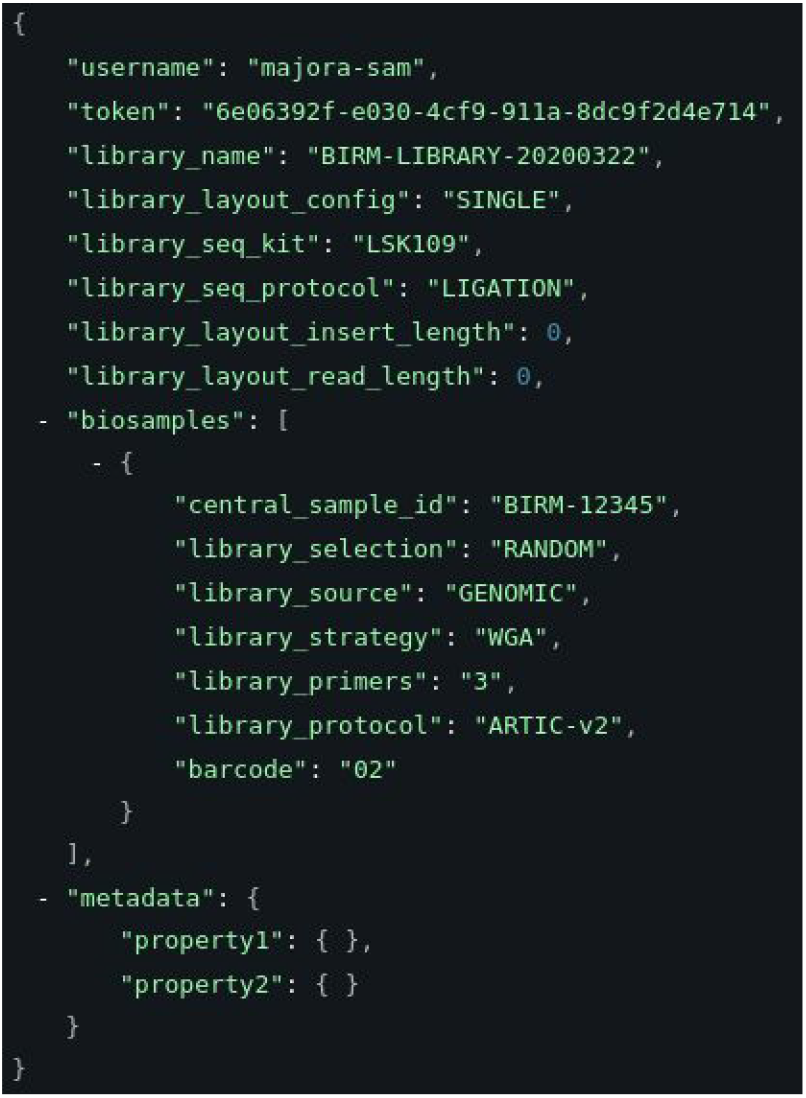
Example API request to submit a new sequencing library artifact to Majora. All metadata from biological samples, to library pooling processes and sequencing runs are communicated to Majora through the various API endpoints. These interfaces take structured data in the JSON format and process them to be stored in the Majora’s SQL database. This example demonstrates the fields and structure of a request to add a new sequencing library to Majora.

To address this, a web application based on the API was built. Briefly, users upload their filled out CSV to a Javascript-based (Nuxt) web frontend. This application transforms the CSV data into the JSON structure mandated by the API and automatically submits it to Majora for validation. Data is transferred securely as the implementation of the uploader tool and the Majora API only supports secure HTTP (https). The Majora API responds with any validation errors that require the user’s attention, and these are parsed and presented prominently in the uploader web application. Valid metadata is accepted immediately and can be queried by any other member of the consortium with access to Majora.

#### A tiered model for the storage of metadata

There are three layers of data within Majora (Table 2). Firstly, for fast searching and filtering, properties common to all instances of a particular type of artifact (e.g. a biosample, or digital file) are explicitly defined as fields in the database. For example, all files have a name, a size and a hash. Database models are implemented based on decisions for the collection and storage of metadata by the various working groups. It is important to be aware of how much work adding or altering fields is: from the technical implementation, through the API, all the way to communicating these changes to users of the template itself.

At the second level, artifacts are a composition of additional models referred to as Metrics. Metrics are models in their own right and can be arbitrarily attached to any artifact in Majora. Metrics are used to store secondary information that is common enough to warrant easy access but not intrinsic to every artifact of the same type. For example, a FASTQ file is a file, but not every file is a FASTQ. A Metric for FASTQ files could therefore annotate a file artifact with information specific to a FASTQ such as the number of sequences and average quality. Typically, Metrics are generated automatically by the API to annotate files.

Thirdly, at the most basic layer, it is possible to “tag” an artifact with arbitrary key-value pairs. Due to the implementation it is more difficult to identify and collect artifacts based on this level of metadata (as the data is not backed by a real model like the first and second layers), but it allows for incredible flexibility. This tertiary metadata level is designed to store information that is not important enough to be incorporated into the primary model and is too esoteric to be part of a secondary metric. The most important part of this layer is its implementation permits any information to be added to an artifact at any time, without any configuration.

### Harmonisation and continuous integration of uploaded sequence and metadata

#### Elan: Autonomous, scalable, daily data integration of sequences and metadata

With FASTA and BAM files uploaded by sequencing sites to CLIMB, and metadata uploaded by users via the API and stored within Majora, these two streams of data must be paired together, processed and published in some way such that they are available to everybody in the consortium who wishes to perform analysis. Once metadata and sequence have been uploaded for a sample, it waits to be pulled into the “inbound distribution” pipeline, named Elan.

Elan is responsible for querying Majora and the CLIMB file system to find unprocessed samples; ensuring the uploaded files are valid, that the BAM contains only SARS-CoV-2 reads, conducts a quality check of the FASTA sequence and BAM alignments, and copies the files to an organised read-only location on CLIMB. If the Majora APIs are the brain of the COG-UK digital infrastructure, then Elan is the heart (Figure 7). The Elan pipeline is run every day and weekly reports are written based on data submitted by Friday, providing a natural cut-off for consortium members to aim to upload their metadata and sequences by.

**Figure 6.**
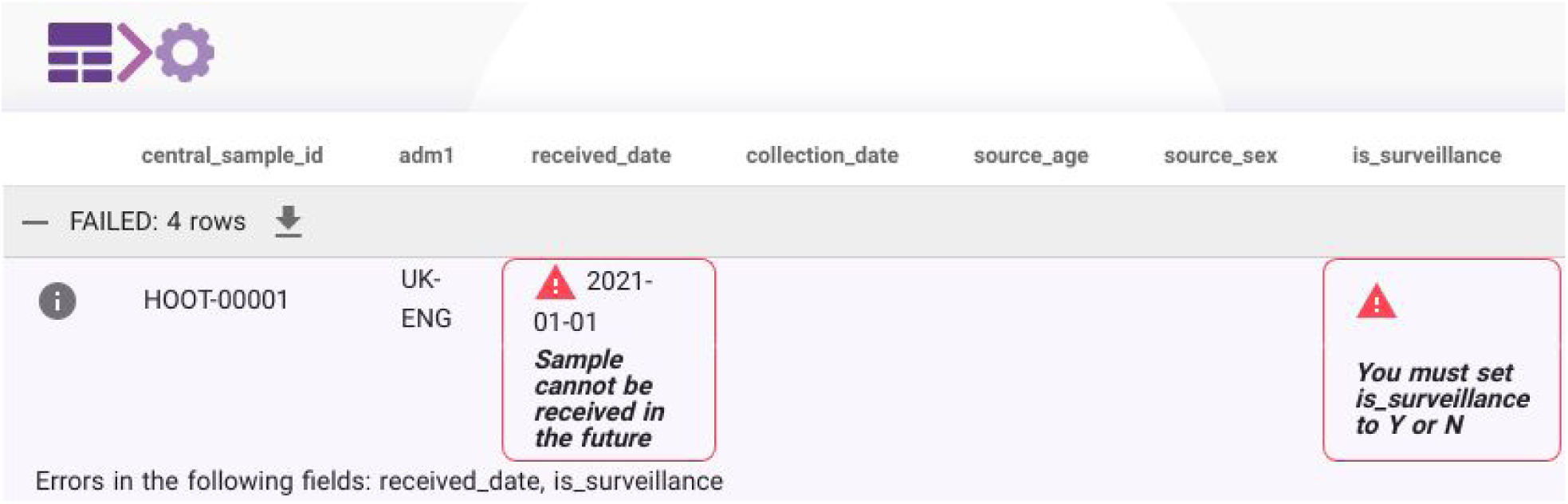
Screenshot of the metadata uploader demonstrating user-facing errors. Metadata is submitted to the consortium by uploading a filled in CSV template to the metadata uploader web application. The uploader converts the CSV data into JSON and communicates with the Majora API. Validation errors are immediately returned, parsed and displayed to the user as shown here.

**Figure 7.**
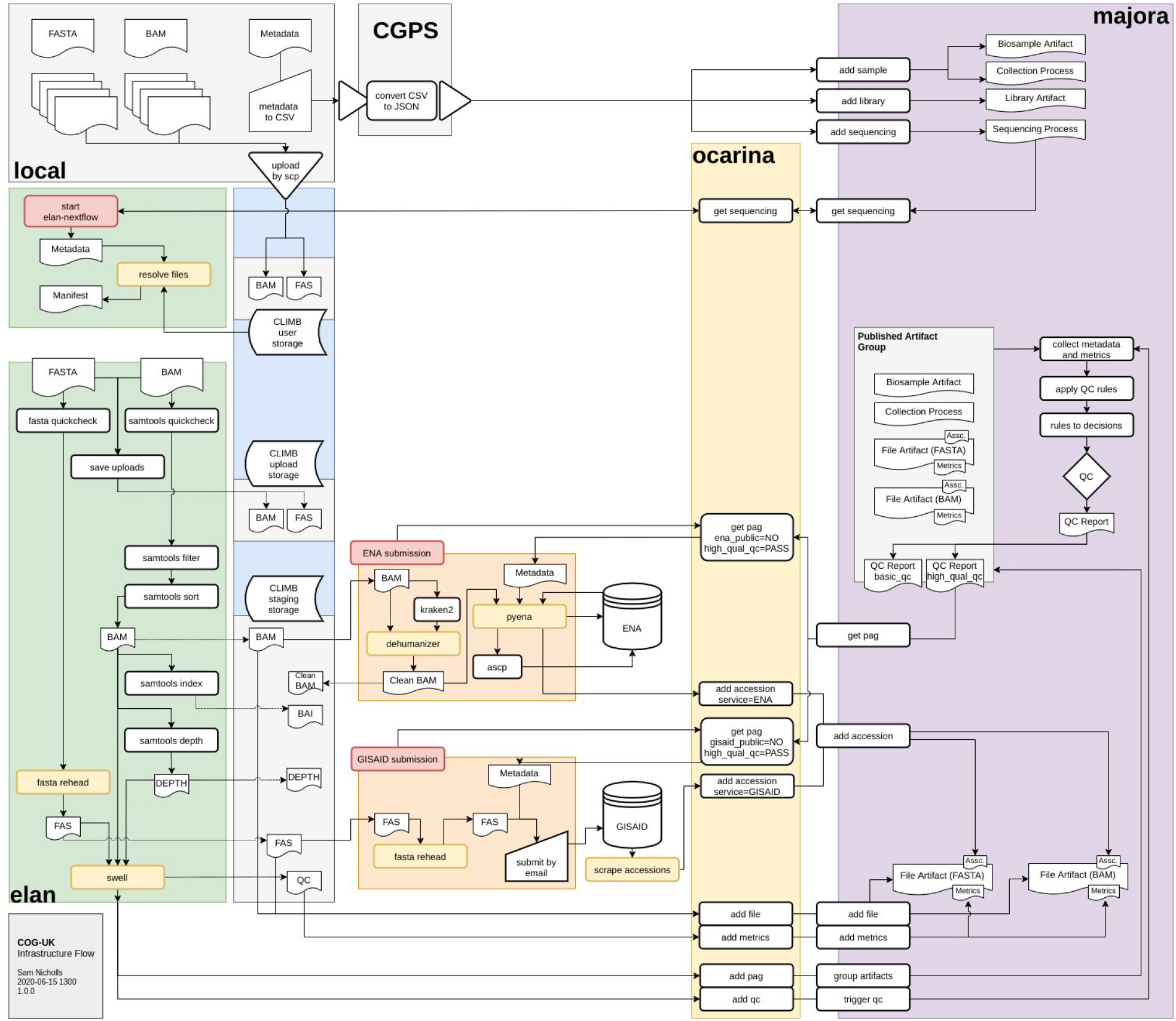
Architecture diagram for the COG-UK Majora platform. Local sites (grey, top left) generate consensus FASTA and alignment BAM for each sequenced sample. Corresponding metadata is collected and managed into a CSV using the consortium template. FASTA and BAM are uploaded to CLIMB using scp or rsync. Metadata is converted from CSV to JSON by the metadata uploader tool and passed to the Majora API to be processed (purple, right). The Elan inbound pipeline (green, left) queries the Majora metadata database using the Ocarina command line client (yellow, right). Elan matches Majora metadata to uploaded files on CLIMB (blue, left) and conducts quality control. Quality metrics are passed to Majora through Ocarina. Downstream pipelines such as outbound distribution pipelines (orange, center) are able to query Majora using Ocarina and package high quality sequences for public databases.

Elan (https://github.com/SamStudio8/elan-nextflow/) is an open-source pipeline built with the NextFlow workflow language [9]. The configuration allows Elan to scale to handle thousands of samples a week by seamlessly executing jobs on additional compute nodes using a job scheduler (SLURM).

#### Orchestrating data flows with human or machine readable messages

Before the daily pipeline is run, Elan uses a web hook to send a series of automated announcements to a well populated Slack channel, notifying users of problems with missing metadata or missing files to fix before the pipeline begins. When Elan finishes, an announcement counting the number of new and cumulative sequences that have passed QC is broadcast (e.g. Figure 8). Elan also emits machine-readable messages (Figure 9) to notify downstream pipelines that there are new samples to process (specifically, we use mqtt [Message Queuing Telemetry Transport], but other similar protocols are available). Using machine-readable messages to control other pipelines reduces human workload and encourages the development of multiple pipelines that do their particular tasks well, rather than tasks being rolled into one monolithic pipeline.

**Figure 8.**
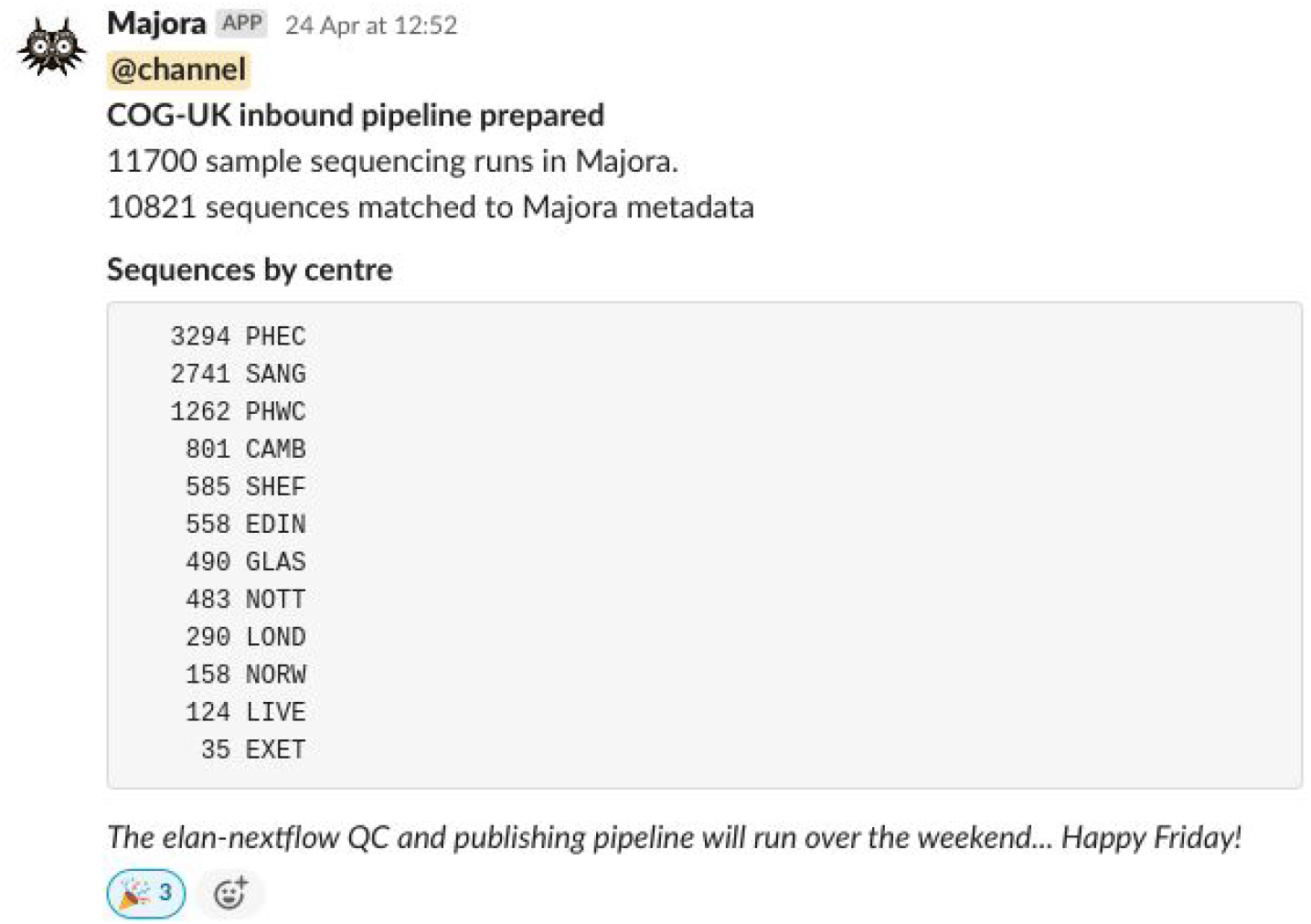
An automated Slack message announcing the start of Elan pipeline. The Elan inbound distribution pipeline is operated transparently by providing a series of courtesy messages before and after it has run. Slack messages are sent programmatically through a web hook to announce samples that appear to be missing metadata or a genome sequence. The example above dated April 24th announces that Elan was about to process the 10,000th sample.

**Figure 9.**
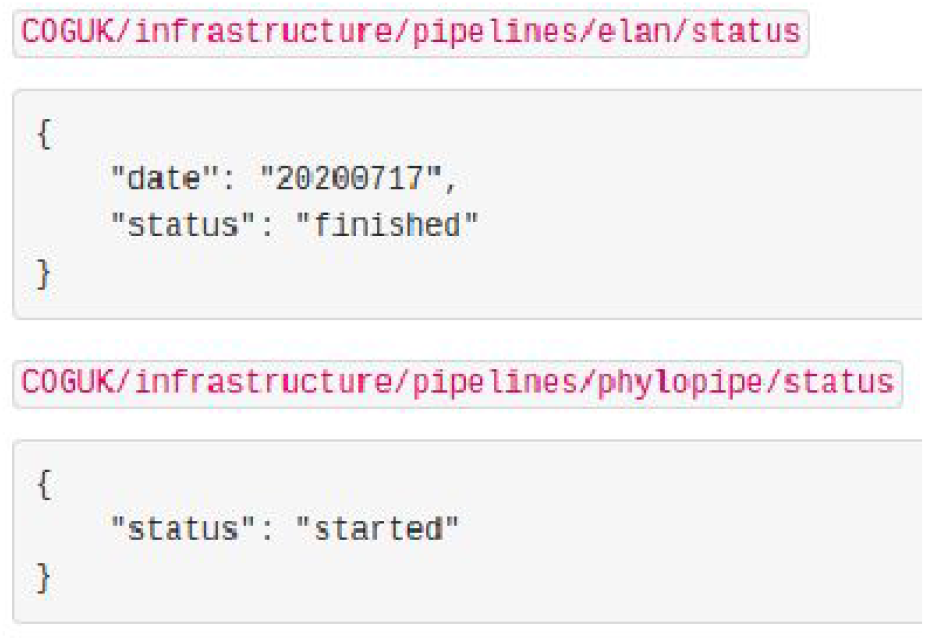
Screenshot of the metadata uploader demonstrating user-facing errors. To assist orchestration of pipelines, we run a message broker service that allows different pipelines within COG-UK to send messages and interact with each other. This example shows Elan emitted a message to announce it has successfully completed, and the phylogenetics pipeline responding to say it has started as a result of the new data to be processed.

#### A QC-aware platform for querying sequences

Majora is also the arbiter of quality control. As Elan processes new sequence FASTA and alignment BAM files, their associated quality metrics are submitted to Majora. Once the metrics are in place, Elan uses an API endpoint to request that Majora carry out a particular QC test and store the result as a QC report.

Quality control tests are composed of rules and decisions based on those rules, and can be imported to Majora by an administrator with a formatted text file. Quality control tests can also be grouped together such that only a subset of tests in the same group are run, with the decision to execute a test or not based on the contents of the sample or sequencing metadata. For example, a QC group may have distinct tests for both Illumina and ONT platforms, but Majora can choose which test to run based on the sequencing metadata submitted by the user (Figure 10).

**Figure 10.**
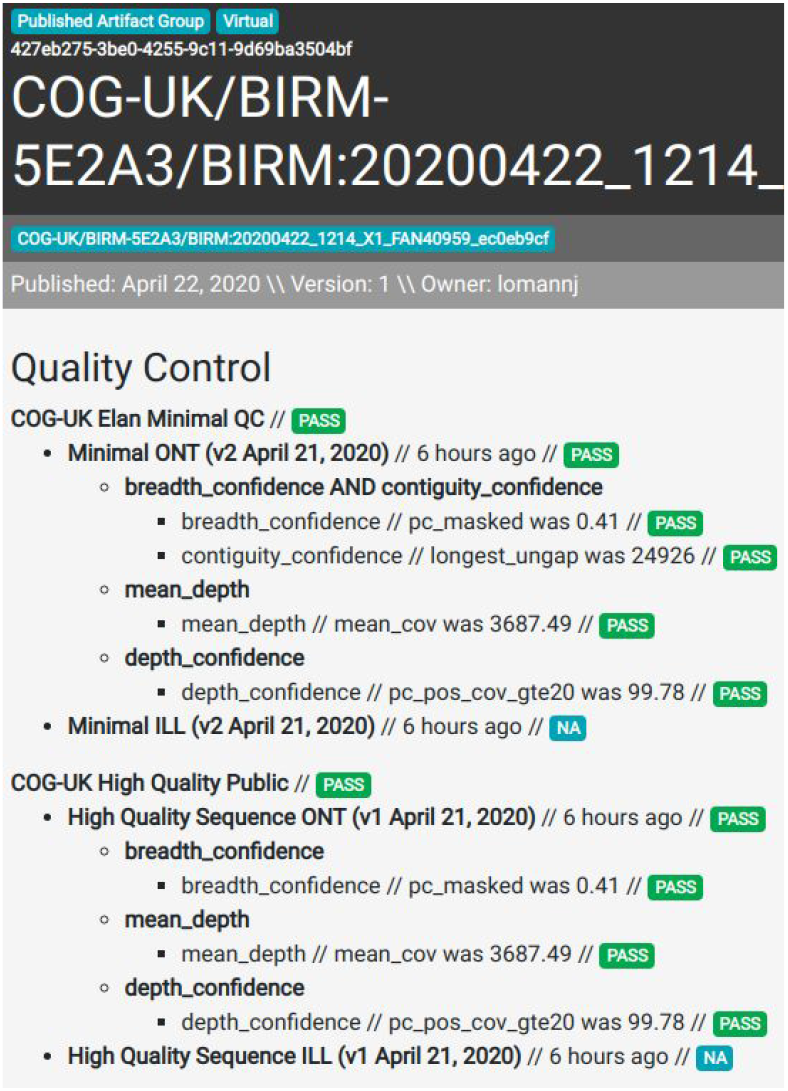
A QC report as viewed through the Majora web application. Quality report view for a real sequence uploaded by a COG-UK site. As part of COG-UK there are two core quality reports generated for every sample. Minimal QC ensures the sample passed basic quality control and is eligible for inclusion in the consortium data set, and the more stringent High Quality report determines whether the sequence will be shared in public databases. Note in this screenshot that each of the two reports have an “ONT” (Oxford Nanopore) and “ILL” (Illumina) entry. In this case, the Illumina tests are skipped because Majora allows tests to be applied based on metadata about the sample. Here, the Illumina tests are skipped because the sequencing metadata shows this run was completed on an Oxford Nanopore sequencer. The two forms of the same test are considered to be part of the same quality report group, which means that the test result is treated equally.

As Majora is aware of the QC reports for each submitted sample, the API endpoints that retrieve data can filter for samples that have passed (or failed) a particular QC test. For the Majora instance deployed on CLIMB-COVID we have two QC test groups: Basic QC is a highly tolerant test which must be passed in order for a sequence to be submitted for downstream pipelines within the consortium, and High Quality QC has a stricter threshold to determine whether sequences will be deposited to public databases. Storing all the quality metrics (e.g. sample cycle threshold value) also allows large-scale comparisons of quality control to be performed across all the consortium’s different sequencing sites and platforms.

#### Routine analysis of the unified data set

When each daily Elan run completes, a machine-readable signal is emitted that initiates the phylogenetics pipeline. This pipeline combines the complete set of non-UK SARS-CoV-2 sequences from GISAID - updated weekly - with the complete set of COG-UK sequences that have passed basic quality control, with the aim of building a phylogenetic tree that captures the evolutionary relationships between all of the sampled viruses to date. The phylogenetics pipeline is written in the Snakemake workflow language [10] and executes a mixture of custom and established software (https://github.com/COG-UK/grapevine).

The pipeline automatically generates reports that summarize the spatial, temporal and genetic diversity of SARS-CoV-2 viruses circulating in the United Kingdom. These reports and other outputs are made available to the consortium through Slack. The global tree and associated metadata also provide the data underlying a bespoke cluster investigation tool which allows users to query sequences of interest and generate reports summarising their phylogenetic and epidemiological context, as well as a publicly available visualisation tool.

#### Linking and visualising consortium data with Microreact

Microreact is a web application that facilitates interpretation of biological data by presenting linked data in multiple panels within a single interactive view [11]. For the COG-UK project five panels are available. 1) A map view showing the place of sample collection; 2) A graph view showing the frequency of lineages over time; 3) A phylogeny derived from the analysis described in the section above; 4) A timeline showing distribution of samples over time; 5) A metadata table view. These views are generated with COG-UK data that has been processed by Elan and the phylogenetics pipeline.

Coarse location metadata from CLIMB is cleaned and geocoded by analysts, and locations are linked to coloured labels with Data-flo (https://data-flo.io). Data-flo provides the ability to manipulate data programmatically and reproducibly using declarative data flows consisting of modular adaptors that perform discrete steps in the overall transformation. The location metadata is combined with the newick phylogeny from the phylogenetics pipeline to output the COG-UK Microreact instance (Figure 11), which includes both the COG-UK data and worldwide data from GISAID (https://microreact.org/project/cogconsortium).

**Figure 11.**
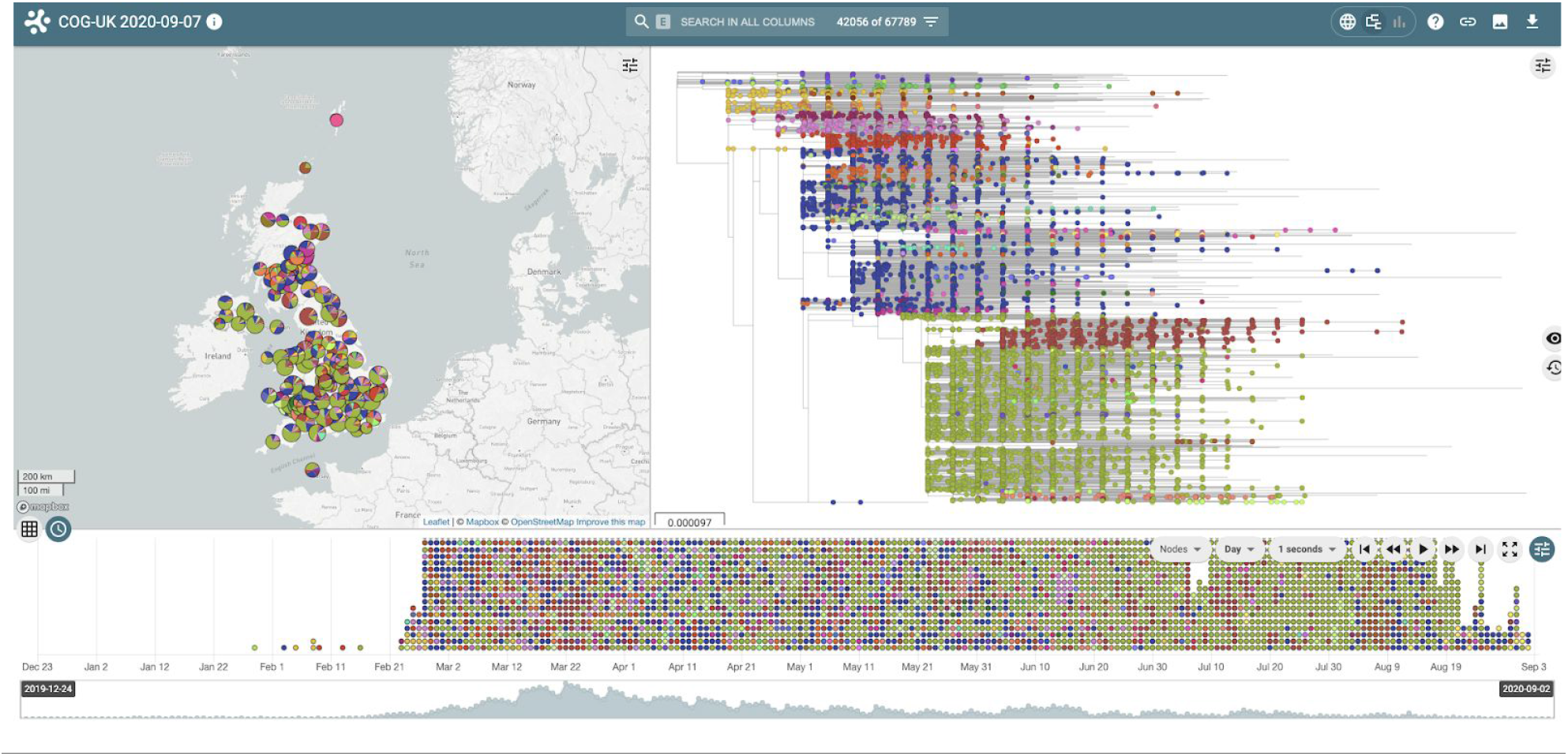
Screenshot of the COG-UK Microreact instance.

The tree viewer is capable of scalable rendering of hundreds of thousands of leaves. This enables full interactivity between all panels such that selecting a subset of samples such as a region on the map, a subtree in the phylogeny or a window within the timeline is instantly reflected in all other panels. This enables querying the data in a visual way that can help inform public health intervention and scientific hypothesis generation. For example selecting a monophyletic group of genetically very similar samples will update the map and timeline and demonstrate if these samples are co-located in time and space and therefore represent a putative outbreak or transmission chain.

#### Controlling access to metadata with view-based permissions

Sample metadata submitted to Majora is divided into one of three access control levels: public, consortium and restricted (Table 1). Most - but not all - public and consortium level data can be viewed through the Majora web interfaces and API. However, when it came to sharing this metadata downstream, we realised that combinations of these fields can in themselves produce a data set that requires a different level of access. It was apparent that granting a user access to a restricted field did not necessarily imply they should have access to all restricted fields (or indeed, all other fields), so it is not enough to simply associate a user with a single access control level.

Rather than granting a user permission to a particular access level, or deploying a cumbersome case-by-case field-level permission system, we control access to metadata by predefining a set of named views that explicitly enumerate a subset of the metadata fields. The view itself then acts as a permission, with users making a case for why they should be granted permission to that view. Access to these views is audited; we always know who can view what data, and when they have used that permission to download a copy of data through a view.

This has several benefits: it is trivial to report who has access to what metadata and provides a common set of named views that can be discussed and shared by downstream analysts. The data view concept also allows us to adhere to the requirements of four different UK public health agencies, each of which acts as a gatekeeper for restricted data from their nation.

The views are implemented in Majora, with configurations loaded by an administrator for a text file. The serializer (which converts the model representation from the database into JSON) is then configured based on the view that has been requested - ensuring that only fields in the view are serialized and sent to the user. Additionally, Majora allows filters to be dynamically applied to the data view to produce derivative data sets. For example, the mechanism through which we share restricted local identifiers to public health agencies will filter based on the country the sample was collected in, and whether a user has signed a particular agreement though Majora. This means a human doesn’t have to spend time sanitising data sets to be distributed.

#### Distributing sequence and metadata outside the consortium

An important goal for the consortium is to provide other projects and scientists outside of COG-UK access to the sequences and limited metadata to be able to perform analysis of their own. This poses another interoperability problem, as sequences and metadata must be converted into a format acceptable by external databases in order to be deposited. COG-UK has made a decision to share data in multiple ways, to ensure that different user groups are able to access the data generated in the simplest way possible.

The Global Initiative on Sharing All Influenza Data (GISAID, [12]) is an established infrastructure for the rapid sharing of sequence data for Influenza. As existing Influenza-focused public health laboratory networks were amongst the first to pivot to sequencing SARS-CoV-2 as part of the pandemic response, GISAID rapidly introduced a capacity for sharing SARS-CoV-2 genomes via its platform. The pre-existing usage of GISAID amongst public health laboratories meant that it quickly gained traction as the *de facto* database to deposit SARS-CoV-2 sequence data. There has been some debate in the wider scientific community about the openness of GISAID and the rules around data access and use, however, within the global health community GISAID is a trusted route for sharing data, which is designed to overcome previous issues of poor academic behavior, particularly in relation to data generated by LMICs. Although there are plans for GISAID depositions to eventually move toward a more automated API-based system, currently the data must be arranged according to their CSV-based template. We have written a command line client capable of requesting particular columns of metadata from Majora to automatically generate a suitable CSV and corresponding FASTA file for weekly submissions to the GISAID database.

To support the objectives of Open Science, as well as meeting the obligations of the FAIR principles (https://www.go-fair.org/fair-principles/) we chose to archive the raw sequencing reads in the European Nucleotide Archive (ENA), which makes the data available internationally through the International Nucleotide Sequence Database Collaboration (INSDC). Making raw reads available is an important step for external researchers to be able to corroborate findings as well as analyze properties of the reads that are lost when only the consensus genomes are available. Our ENA submission pipeline takes care to mitigate the risk of inadvertently sharing human data (https://github.com/SamStudio8/dehumanizer).

Automated submissions to GISAID and the ENA are provided as a service to members of the consortium. Site leads can opt their institutes into automated submissions from the Majora website. Accessions are shared within and outside the consortium. For raw reads, our pyENA command line client captures accessions as part of the submission protocol. For GISAID, some manual intervention is required as these cannot be extracted programmatically.

## Discussion and Conclusion

In this article we have described the end-to-end compute infrastructure we developed for the COVID-19 Genomics UK (COG-UK) consortium. Our platform addresses the needs of a distributed democratised network for sequencing SARS-CoV-2 genomes, providing a unified interface for transferring, storing and sharing sequence and metadata. The system provides a core platform for harmonisation and continuous integration of uploaded sequence and metadata which has underpinned the activities of COG-UK, analysing over 50,000 SARS-CoV-2 genomes since its inception.

An agile approach to development allowed us to respond quickly to changes in the needs of the project. This was especially important given the diverse array of wet and dry laboratory protocols used across all the different members of the consortium. COG-UK’s success is owed in part to its agility to turn around a response to a novel pathogen. It would be fair to describe the development process as “reactive”. We did not set out to build a perfect system from the beginning, allowing the constraints we encountered to guide design decisions as and when they needed to be made.

Hosting this infrastructure on CLIMB is both a pragmatic and perfect choice. CLIMB is probably still the largest dedicated compute infrastructure for microbial genomics in the world. The shared nature of the platform was critical for immediate sharing and analysis. Within three days of booting the first virtual machine we were receiving uploads of sequence data. Within a week, 260 complete genomes from 7 sequencing centres had been uploaded and processed by our inbound distribution pipeline - already more genomes than any other country in the world other than China at the time. Within two months, COG-UK was responsible for half of all the international SARS-CoV-2 sequences deposited into GISAID.

Although Black et al. [6] recently suggested it “would be easier to licence databasing software for the metadata database than to build it from scratch”, we had the expertise in place to rapidly develop appropriate software that was unlike anything on the market. Architecting our own database has allowed the metadata definitions, metadata templates and database to evolve together with the changing demands of the consortium. Our development methodology focussed on building a minimal viable product to address the current needs of the consortium. Although our agile development allowed us to move quickly, it does not compromise on functionality: our platform has been built from the ground up by people with domain knowledge, given the opportunity to start over, we would make many of the same design choices again.

The infrastructure we have built from scratch covers application programming interfaces (APIs) for uploading metadata and sequence data to an isolated server on CLIMB, automated pipelines for quality control and organising data within the consortium, processes to automatically share data outside the consortium and the core infrastructure for the secure storage and management of data assets on the system.

The success of this system has depended on a close working relationship between analysis teams, sample laboratories, the template authors, the authors of the uploading tool and the author of the Majora API.

Along with the technical pragmatism we have described in this article, future responses should aim to generate better identifiers and attempt to simplify data sharing (public health agencies could generate specific “random” identifiers for a record and share them) or invest in a secure salting scheme. The availability of single, unique, shareable identifiers across a geographically and organisationally dispersed consortium has been one of the largest obstacles to the work of the consortium. With so many NHS organisations composing the national service for each of the countries in the UK, with the exception of Wales, there is not a single centralised organisation that can mandate data sharing.

Our difficulties in obtaining sample and patient identifiers has made it more difficult to collate multiple samples from the same individual and delays in security assessments and contractual arrangements for using granular geographic data left analysts with the unfortunate task of munging various different representations of counties and cities within the United Kingdom. These metadata issues highlight a need for future readiness, not just for technical solutions, but regulatory ones too. To be ready for the next pandemic, we must be able to generate suitable sample and patient identifiers, we must simplify access to minimally useful metadata, and enact a regulatory framework to share identifiers and core information like dates and postcodes in an effective manner.

The infrastructure we have presented here is generalizable to future novel pathogens, but could also be expanded to cover metagenomics and environmental sampling. CLIMB-COVID is a proven model, evidenced by the success of the COG-UK consortium (Figure 12). As of writing, COG-UK has produced 50,000 public sequences, has contributed more than 10 reports to the government, supported more than 50 outbreak investigations in the UK and has been used in >16 publications.

**Figure 12.**
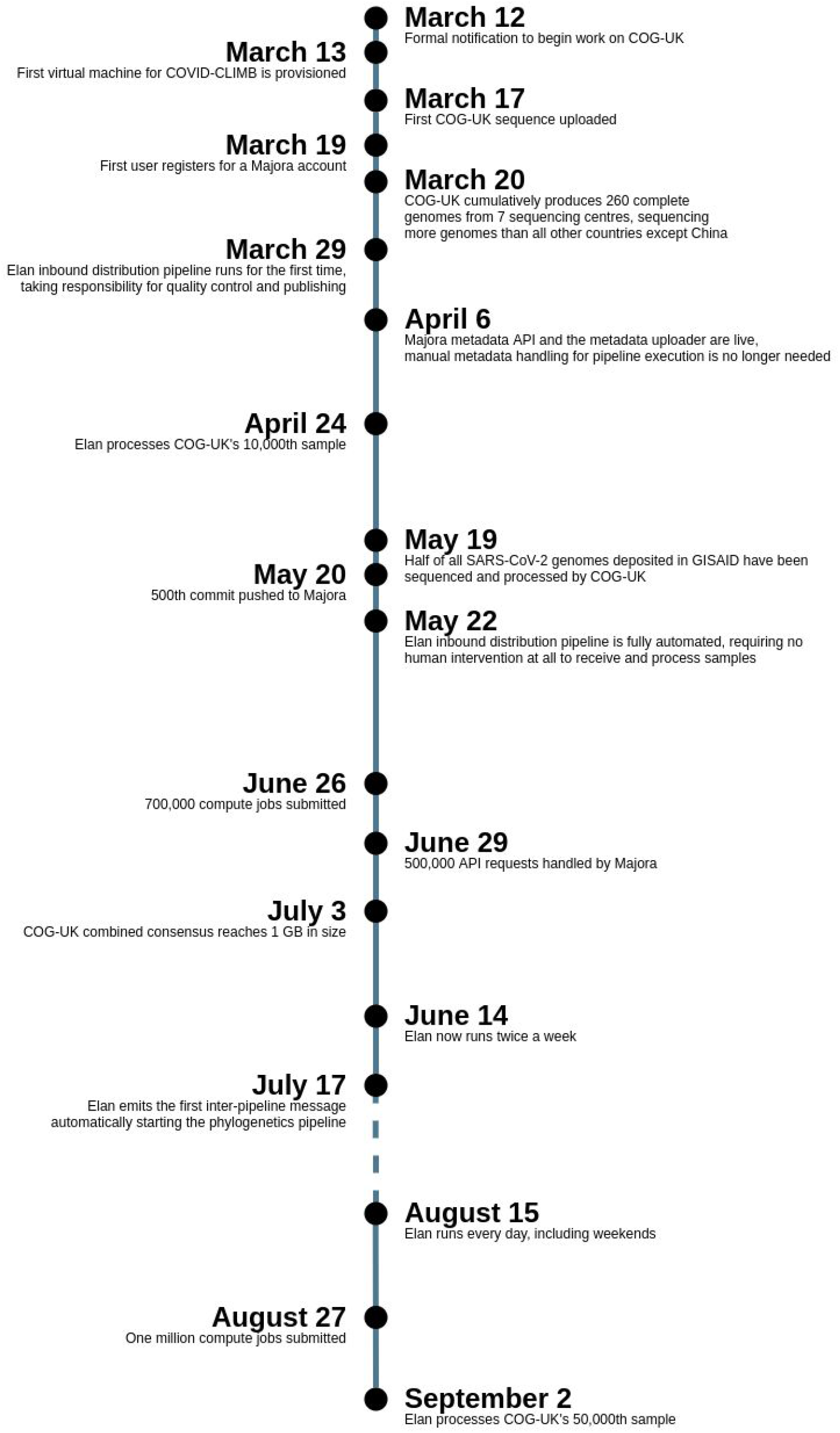
COG-UK platform timeline and performance milestones.

Our efforts have enabled us to go from a blank slate to an integrated infrastructure that colaesces the sequence and metadata from multiple sequencing centres spread across four distinct healthcare systems. In normal times this would be considered a considerable success - to do it in the middle of a pandemic is extraordinary. The template we present here should therefore be an example for those who have similar objectives, as well as presenting a very different vision to those who would suggest that data should be centralised into databases that sit apart from analysis tools and detailed medata.

## Data availability

Majora is a Django web application for tracking artifacts and processes, it is open source software available via github.com/SamStudio8/majora. The Ocarina command line client reference implementation for using the Majora API is open source and freely distributed under the MIT license via github.com/SamStudio8/ocarina. The Elan inbound distribution Nextflow pipeline is open source and freely distributed under the MIT license via github.com/SamStudio8/elan-nextflow. The Swell program to calculate QC metrics from BAM depth files is open source and freely distributed under the MIT license via github.com/SamStudio8/swell. The Dehumanizer program to sanitize BAMs is open source and freely distributed under the MIT license via github.com/SamStudio8/dehumanizer. The PyENA program to upload BAMs to ENA is open source and freely distributed under the MIT license via github.com/SamStudio8/pyena.

The Grapevine phylogenetics Snakemake pipeline is open source and available for download via github.com/COG-UK/grapevine. The ARTIC data processing Nextflow pipeline is open source and available for download via github.com/connor-lab/ncov2019-artic-nf.

Consensus SARS-CoV-2 genomes are routinely deposited into GISAID and also made available via cogconsortium.uk/data. COG-UK data can be explored using a Microreact instance available at microreact.org/project/cogconsortium. Human-filtered sequencing data for COG-UK are routinely deposited in the European Nucleotide Archive (ENA) at EMBL-EBI under accession PRJEB37886.

## Funding and acknowledgements

COG-UK is supported by funding from the Medical Research Council (MRC) part of UK Research & Innovation (UKRI), the National Institute of Health Research (NIHR) and Genome Research Limited, operating as the Wellcome Sanger Institute. CLIMB is funded by the Medical Research Council (MRC) through grant MR/L015080/1. The authors would like to thank the Birmingham Environment for Academic Research (BEAR) at the University of Birmingham for supporting the deployment of this project.

## Author contributions

Conceptualization NL SN RP MB

Data curation SN

Formal Analysis SN

Funding acquisition SP NL TC DA AR

Investigation SN

Methodology SN MC

Project administration EH NL TC SG DA AR

Resources SN RP MB SC RL COG-UK

Software SN RP MB AU KD BT BJ SR RA

Supervision NL

Validation SN

Visualization SN

Writing – original draft SN

Writing – review & editing SN NL TC AU BJ MC DA

## Supplementary information

Full COG-UK metadata specification

### The COVID-19 Genomics UK (COG-UK) Consortium

**Funding acquisition, leadership, supervision, metadata curation, project administration, samples, logistics, Sequencing, analysis, and Software and analysis tools:**

Thomas R Connor ^33, 34^, and Nicholas J Loman ^15^.

**Leadership, supervision, sequencing, analysis, funding acquisition, metadata curation, project administration, samples, logistics, and visualisation:**

Samuel C Robson ^68^.

**Leadership, supervision, project administration, visualisation, samples, logistics, metadata curation and software and analysis tools:**

Tanya Golubchik ^27^.

**Leadership, supervision, metadata curation, project administration, samples, logistics sequencing and analysis:**

M. Estee Torok ^8, 10^.

**Project administration, metadata curation, samples, logistics, sequencing, analysis, and software and analysis tools**

William L Hamilton ^8, 10^.

**Leadership, supervision, samples logistics, project administration, funding acquisition sequencing and analysis:**

David Bonsall ^27^.

**Leadership and supervision, sequencing, analysis, funding acquisition, visualisation and software and analysis tools:**

Ali R Awan ^74^.

**Leadership and supervision, funding acquisition, sequencing, analysis, metadata curation, samples and logistics:**

Sally Corden^33^.

**Leadership supervision, sequencing analysis, samples, logistics, and metadata curation:**

Ian Goodfellow ^11^.

**Leadership, supervision, sequencing, analysis, samples, logistics, and Project administration:**

Darren L Smith ^60, 61^.

**Project administration, metadata curation, samples, logistics, sequencing and analysis:**

Martin D Curran ^14^, and Surendra Parmar ^14^.

**Samples, logistics, metadata curation, project administration sequencing and analysis:**

James G Shepherd ^21^.

**Sequencing, analysis, project administration, metadata curation and software and analysis tools:**

Matthew D Parker ^38^.

**Leadership, supervision, funding acquisition, samples, logistics, and metadata curation:**

Catherine Moore ^33^.

**Leadership, supervision, metadata curation, samples, logistics, sequencing and analysis:**

Derek J Fairley^6, 88^, Matthew W Loose ^54^, and Joanne Watkins ^33^.

**Metadata curation, sequencing, analysis, leadership, supervision and software and analysis tools:**

Matthew Bull ^33^, and Sam Nicholls ^15^.

**Leadership, supervision, visualisation, sequencing, analysis and software and analysis tools:**

David M Aanensen ^1, 30^.

**Sequencing, analysis, samples, logistics, metadata curation, and visualisation:**

Sharon Glaysher ^70^.

**Metadata curation, sequencing, analysis, visualisation, software and analysis tools:**

Matthew Bashton ^60^, and Nicole Pacchiarini ^33^.

**Sequencing, analysis, visualisation, metadata curation, and software and analysis tools:**

Anthony P Underwood ^1, 30^.

**Funding acquisition, leadership, supervision and project administration:**

Thushan I de Silva ^38^, and Dennis Wang ^38^.

**Project administration, samples, logistics, leadership and supervision**:

Monique Andersson^28^, Anoop J Chauhan ^70^, Mariateresa de Cesare ^26^, Catherine Ludden ^1,3^, and Tabitha W Mahungu^91^.

**Sequencing, analysis, project administration and metadata curation:**

Rebecca Dewar ^20^, and Martin P McHugh ^20^.

**Samples, logistics, metadata curation and project administration:**

Natasha G Jesudason ^21^, Kathy K Li MBBCh ^21^, Rajiv N Shah ^21^, and Yusri Taha ^66^.

**Leadership, supervision, funding acquisition and metadata curation:**

Kate E Templeton ^20^.

**Leadership, supervision, funding acquisition, sequencing and analysis:**

Simon Cottrell ^33^, Justin O’Grady ^51^, Andrew Rambaut ^19^, and Colin P Smith^93^.

**Leadership, supervision, metadata curation, sequencing and analysis:**

Matthew T.G. Holden ^87^, and Emma C Thomson ^21^.

**Leadership, supervision, samples, logistics and metadata curation**:

Samuel Moses ^81, 82^.

**Sequencing, analysis, leadership, supervision, samples and logistics:**

Meera Chand ^7^, Chrystala Constantinidou ^71^, Alistair C Darby ^46^, Julian A Hiscox ^46^, Steve Paterson ^46^, and Meera Unnikrishnan ^71^.

**Sequencing, analysis, leadership and supervision and software and analysis tools:**

Andrew J Page ^51^, and Erik M Volz ^96^.

**Samples, logistics, sequencing, analysis and metadata curation:**

Charlotte J Houldcroft ^8^, Aminu S Jahun ^11^, James P McKenna ^88^, Luke W Meredith ^11^, Andrew Nelson ^61^, Sarojini Pandey ^72^, and Gregory R Young ^60^.

**Sequencing, analysis, metadata curation, and software and analysis tools:**

Anna Price ^34^, Sara Rey ^33^, Sunando Roy ^41^, Ben Temperton^49^, and Matthew Wyles ^38^.

**Sequencing, analysis, metadata curation and visualisation:**

Stefan Rooke^19^, and Sharif Shaaban ^87^.

**Visualisation, sequencing, analysis and software and analysis tools:**

Helen Adams ^35^, Yann Bourgeois ^69^, Katie F Loveson ^68^, Áine O’Toole ^19^, and Richard Stark ^71^.

**Project administration, leadership and supervision:**

Ewan M Harrison ^1, 3^, David Heyburn ^33^, and Sharon J Peacock ^2, 3^

**Project administration and funding acquisition:**

David Buck ^26^, and Michaela John^36^

**Sequencing, analysis and project administration:**

Dorota Jamrozy ^1^, and Joshua Quick ^15^

**Samples, logistics, and project administration:**

Rahul Batra ^78^, Katherine L Bellis ^1, 3^, Beth Blane ^3^, Sophia T Girgis ^3^, Angie Green ^26^, Anita Justice ^28^, Mark Kristiansen ^41^, and Rachel J Williams ^41^.

**Project administration, software and analysis tools:**

Radoslaw Poplawski^15^.

**Project administration and visualisation:**

Garry P Scarlett ^69^.

**Leadership, supervision, and funding acquisition:**

John A Todd ^26^, Christophe Fraser ^27^, Judith Breuer ^40,41^, Sergi Castellano ^41^, Stephen L Michell ^49^, Dimitris Gramatopoulos ^73^, and Jonathan Edgeworth ^78^.

**Leadership, supervision and metadata curation:**

Gemma L Kay ^51^.

**Leadership, supervision, sequencing and analysis:**

Ana da Silva Filipe ^21^, Aaron R Jeffries ^49^, Sascha Ott ^71^, Oliver Pybus ^24^, David L Robertson ^21^, David A Simpson ^6^, and Chris Williams ^33^.

**Samples, logistics, leadership and supervision:**

Cressida Auckland ^50^, John Boyes ^83^, Samir Dervisevic ^52^, Sian Ellard ^49, 50^, Sonia Goncalves^1^, Emma J Meader ^51^, Peter Muir ^2^, Husam Osman ^95^, Reenesh Prakash ^52^, Venkat Sivaprakasam ^18^, and Ian B Vipond ^2^.

**Leadership, supervision and visualisation**

Jane AH Masoli ^49, 50^.

**Sequencing, analysis and metadata curation**

Nabil-Fareed Alikhan ^51^, Matthew Carlile ^54^, Noel Craine ^33^, Sam T Haldenby ^46^, Nadine Holmes ^54^, Ronan A Lyons ^37^, Christopher Moore ^54^, Malorie Perry ^33^, Ben Warne ^80^, and Thomas Williams ^19^.

**Samples, logistics and metadata curation:**

Lisa Berry ^72^, Andrew Bosworth ^95^, Julianne Rose Brown ^40^, Sharon Campbell ^67^, Anna Casey ^17^, Gemma Clark ^56^, Jennifer Collins ^66^, Alison Cox ^43, 44^, Thomas Davis ^84^, Gary Eltringham ^66^, Cariad Evans ^38, 39^, Clive Graham ^64^, Fenella Halstead ^18^, Kathryn Ann Harris ^40^, Christopher Holmes ^58^, Stephanie Hutchings ^2^, Miren Iturriza-Gomara ^46^, Kate Johnson ^38, 39^, Katie Jones ^72^, Alexander J Keeley ^38^, Bridget A Knight ^49, 50^, Cherian Koshy^90^, Steven Liggett ^63^, Hannah Lowe ^81^, Anita O Lucaci ^46^, Jessica Lynch ^25, 29^, Patrick C McClure ^55^, Nathan Moore ^31^, Matilde Mori ^25, 29, 32^, David G Partridge ^38, 39^, Pinglawathee Madona ^43, 44^, Hannah M Pymont ^2^, Paul Anthony Randell ^43, 44^, Mohammad Raza ^38, 39^, Felicity Ryan ^81^, Robert Shaw ^28^, Tim J Sloan ^57^, and Emma Swindells ^65^.

**Sequencing, analysis, Samples and logistics:**

Alexander Adams ^33^, Hibo Asad ^33^, Alec Birchley ^33^, Tony Thomas Brooks ^41^, Giselda Bucca ^93^, Ethan Butcher ^70^, Sarah L Caddy ^13^, Laura G Caller ^2, 3, 12^, Yasmin Chaudhry ^11^, Jason Coombes ^33^, Michelle Cronin ^33^, Patricia L Dyal ^41^, Johnathan M Evans ^33^, Laia Fina ^33^, Bree Gatica-Wilcox ^33^, Iliana Georgana ^11^, Lauren Gilbert ^33^, Lee Graham ^33^, Danielle C Groves ^38^, Grant Hall ^11^, Ember Hilvers ^33^, Myra Hosmillo ^11^, Hannah Jones ^33^, Sophie Jones ^33^, Fahad A Khokhar ^13^, Sara Kumziene-Summerhayes ^33^, George MacIntyre-Cockett ^26^, Rocio T Martinez Nunez ^94^, Caoimhe McKerr ^33^, Claire McMurray ^15^, Richard Myers ^7^, Yasmin Nicole Panchbhaya ^41^, Malte L Pinckert ^11^, Amy Plimmer ^33^, Joanne Stockton ^15^, Sarah Taylor ^33^, Alicia Thornton ^7^, Amy Trebes ^26^, Alexander J Trotter ^51^, Helena Jane Tutill ^41^, Charlotte A Williams ^41^, Anna Yakovleva ^11^ and Wen C Yew ^62^.

**Sequencing, analysis and software and analysis tools:**

Mohammad T Alam ^71^, Laura Baxter ^71^, Olivia Boyd ^96^, Fabricia F. Nascimento ^96^, Timothy M Freeman ^38^, Lily Geidelberg ^96^, Joseph Hughes ^21^, David Jorgensen ^96^, Benjamin B Lindsey ^38^, Richard J Orton ^21^, Manon Ragonnet-Cronin ^96^ Joel Southgate ^33, 34,^ and Sreenu Vattipally ^21^.

**Samples, logistics and software and analysis tools:**

Igor Starinskij ^23^.

**Visualisation and software and analysis tools:**

Joshua B Singer ^21^, Khalil Abudahab ^1, 30^, Leonardo de Oliveira Martins ^51^, Thanh Le-Viet ^51^, Mirko Menegazzo ^30^, Ben EW Taylor ^1, 30^, and Corin A Yeats ^30^.

**Project Administration:**

Sophie Palmer ^3^, Carol M Churcher ^3^, Alisha Davies ^33^, Elen De Lacy ^33^, Fatima Downing ^33^, Sue Edwards ^33^, Nikki Smith ^38^, Francesc Coll ^97^, Nazreen F Hadjirin ^3^ and Frances Bolt ^44, 45^.

**Leadership and supervision:**

Alex Alderton^1^, Matt Berriman^1^, Ian G Charles ^51^, Nicholas Cortes ^31^, Tanya Curran ^88^, John Danesh^1^, Sahar Eldirdiri ^84^, Ngozi Elumogo ^52^, Andrew Hattersley ^49, 50^, Alison Holmes ^44, 45^, Robin Howe ^33^, Rachel Jones ^33^, Anita Kenyon ^84^, Robert A Kingsley ^51^, Dominic Kwiatkowski ^1, 9^, Cordelia Langford^1^, Jenifer Mason^48^, Alison E Mather ^51^, Lizzie Meadows ^51^, Sian Morgan ^36^, James Price ^44, 45^, Trevor I Robinson ^48^, Giri Shankar ^33^, John Wain ^51^, and Mark A Webber ^51^.

**Metadata curation:**

Declan T Bradley ^5, 6^, Michael R Chapman ^1, 3, 4^, Derrick Crooke ^28^, David Eyre ^28^, Martyn Guest ^34^, Huw Gulliver ^34^, Sarah Hoosdally ^28^, Christine Kitchen ^34^, Ian Merrick ^34^, Siddharth Mookerjee ^44, 45^, Robert Munn ^34^, Timothy Peto ^28^, Will Potter ^52^, Dheeraj K Sethi ^52^, Wendy Smith ^56^, Luke B Snell ^75, 94^, Rachael Stanley ^52^, Claire Stuart ^52^ and Elizabeth Wastenge^20^.

**Sequencing and analysis:**

Erwan Acheson ^6^, Safiah Afifi ^36^, Elias Allara ^2, 3^, Roberto Amato ^1^, Adrienn Angyal ^38^, Elihu Aranday-Cortes ^21^, Cristina Ariani ^1^, Jordan Ashworth ^19^, Stephen Attwood ^24^, Alp Aydin ^51^, David J Baker ^51^, Carlos E Balcazar ^19^, Angela Beckett ^68^ Robert Beer ^36^, Gilberto Betancor ^76^, Emma Betteridge ^1^, David Bibby ^7^, Daniel Bradshaw^7^, Catherine Bresner ^34^, Hannah E Bridgewater ^71^, Alice Broos ^21^, Rebecca Brown ^38^, Paul E Brown ^71^, Kirstyn Brunker ^22^, Stephen N Carmichael ^21^, Jeffrey K. J. Cheng ^71^, Dr Rachel Colquhoun ^19^, Gavin Dabrera ^7^, Johnny Debebe ^54^, Eleanor Drury ^1^, Louis du Plessis ^24^, Richard Eccles ^46^, Nicholas Ellaby ^7^, Audrey Farbos ^49^, Ben Farr ^1^, Jacqueline Findlay ^41^, Chloe L Fisher ^74^, Leysa Marie Forrest ^41^, Sarah Francois ^24^, Lucy R. Frost ^71^, William Fuller^34^, Eileen Gallagher ^7^, Michael D Gallagher ^19^, Matthew Gemmell ^46^, Rachel AJ Gilroy ^51^, Scott Goodwin ^1^, Luke R Green ^38^, Richard Gregory ^46^, Natalie Groves ^7^, James W Harrison ^49^, Hassan Hartman ^7^, Andrew R Hesketh ^93^,Verity Hill ^19^, Jonathan Hubb ^7^, Margaret Hughes^46^, David K Jackson ^1^, Ben Jackson ^19^, Keith James ^1^, Natasha Johnson ^21^, Ian Johnston ^1^, Jon-Paul Keatley ^1^, Moritz Kraemer ^24^, Angie Lackenby ^7^, Mara Lawniczak ^1^, David Lee ^7^, Rich Livett ^1^, Stephanie Lo ^1^, Daniel Mair ^21^, Joshua Maksimovic ^36^, Nikos Manesis ^7^, Robin Manley ^49^, Carmen Manso ^7^, Angela Marchbank ^34^, Inigo Martincorena ^1^, Tamyo Mbisa ^7^, Kathryn McCluggage ^36^, JT McCrone ^19^, Shahjahan Miah ^7^, Michelle L Michelsen ^49^, Mari Morgan ^33^, Gaia Nebbia ^78^,Charlotte Nelson ^46^, Jenna Nichols ^21^, Paola Niola ^41^, Kyriaki Nomikou ^21^, Steve Palmer ^1^, Naomi Park ^1^, Yasmin A Parr ^1^, Paul J Parsons ^38^, Vineet Patel ^7^, Minal Patel ^1^, Clare Pearson ^2, 1^, Steven Platt ^7^, Christoph Puethe ^1^, Mike Quail ^1^,Jayna Raghwani ^24^, Lucille Rainbow ^46^, Shavanthi Rajatileka ^1^, Mary Ramsay ^7^, Paola C Resende Silva ^41, 42^, Steven Rudder 51, Chris Ruis ^3^, Christine M Sambles ^49^, Fei Sang ^54^, Ulf Schaefer^7^, Emily Scher ^19^, Carol Scott ^1^, Lesley Shirley ^1^, Adrian W Signell ^76^, John Sillitoe ^1^, Christen Smith ^1^, Dr Katherine L Smollett ^21^, Karla Spellman ^36^, Thomas D Stanton ^19^, David J Studholme ^49^, Grace Taylor-Joyce ^71^, Ana P Tedim ^51^, Thomas Thompson ^6^, Nicholas M Thomson ^51^, Scott Thurston^1^, Lily Tong ^21^, Gerry Tonkin-Hill ^1^, Rachel M Tucker ^38^, Edith E Vamos ^4^, Tetyana Vasylyeva^24^, Joanna Warwick-Dugdale ^49^, Danni Weldon ^1^, Mark Whitehead ^46^, David Williams ^7^, Kathleen A Williamson ^19^,Harry D Wilson ^76^,Trudy Workman ^34^, Muhammad Yasir^51^, Xiaoyu Yu ^19^, and Alex Zarebski ^24^.

**Samples and logistics:**

Evelien M Adriaenssens ^51^, Shazaad S Y Ahmad ^2, 47^, Adela Alcolea-Medina ^59, 77^, John Allan ^60^, Patawee Asamaphan ^21^, Laura Atkinson ^40^, Paul Baker ^63^, Jonathan Ball ^55^, Edward Barton^64^, Mathew A Beale^1^, Charlotte Beaver^1^, Andrew Beggs ^16^, Andrew Bell ^51^, Duncan J Berger ^1^, Louise Berry. ^56^, Claire M Bewshea ^49^, Kelly Bicknell ^70^, Paul Bird ^58^, Chloe Bishop ^7^, Tim Boswell ^56^, Cassie Breen ^48^, Sarah K Buddenborg^1^, Shirelle Burton-Fanning ^66^, Vicki Chalker ^7^, Joseph G Chappell ^55^, Themoula Charalampous ^78, 94^, Claire Cormie^3^, Nick Cortes^29, 25^, Lindsay J Coupland ^52^, Angela Cowell ^48^, Rose K Davidson ^53^, Joana Dias ^3^, Maria Diaz ^51^, Thomas Dibling^1^, Matthew J Dorman^1^, Nichola Duckworth^57^, Scott Elliott^70^, Sarah Essex^63^, Karlie Fallon ^58^, Theresa Feltwell ^8^, Vicki M Fleming ^56^, Sally Forrest ^3^, Luke Foulser^1^, Maria V Garcia-Casado^1^, Artemis Gavriil ^41^, Ryan P George ^47^, Laura Gifford ^33^, Harmeet K Gill ^3^, Jane Greenaway ^65^, Luke Griffith^53^, Ana Victoria Gutierrez^51^, Antony D Hale ^85^, Tanzina Haque ^91^, Katherine L Harper ^85^, Ian Harrison ^7^, Judith Heaney ^89^, Thomas Helmer ^58^, Ellen E Higginson^3^, Richard Hopes ^2^, Hannah C Howson-Wells ^56^, Adam D Hunter ^1^, Robert Impey ^70^, Dianne Irish-Tavares ^91^, David A Jackson^1^, Kathryn A Jackson ^46^, Amelia Joseph ^56^, Leanne Kane ^1^, Sally Kay ^1^, Leanne M Kermack ^3^, Manjinder Khakh ^56^, Stephen P Kidd ^29, 25,31^, Anastasia Kolyva ^51^, Jack CD Lee ^40^, Laura Letchford ^1^, Nick Levene ^79^, Lisa J Levett ^89^, Michelle M Lister ^56^, Allyson Lloyd ^70^, Joshua Loh ^60^, Louissa R Macfarlane-Smith ^85^, Nicholas W Machin ^2, 47^, Mailis Maes ^3^, Samantha McGuigan ^1^, Liz McMinn ^1^, Lamia Mestek-Boukhibar ^41^, Zoltan Molnar ^6^, Lynn Monaghan ^79^, Catrin Moore ^27^, Plamena Naydenova ^3^, Alexandra S Neaverson ^1^, Rachel Nelson ^1^, Marc O Niebel ^21^, Elaine O’Toole^48^, Debra Padgett ^64^, Gaurang Patel ^1^, Brendan AI Payne ^66^, Liam Prestwood ^1^, Veena Raviprakash ^67^, Nicola Reynolds^86^, Alex Richter ^16^, Esther Robinson ^95^, Hazel A Rogers^1^, Aileen Rowan ^96^, Garren Scott ^64^, Divya Shah ^40^, Nicola Sheriff ^67^, Graciela Sluga, Emily Souster^1^, Michael Spencer-Chapman^1^, Sushmita Sridhar ^1, 3^, Tracey Swingler ^53^, Julian Tang^58^, Graham P Taylor^96^, Theocharis Tsoleridis ^55^, Lance Turtle^46^, Sarah Walsh ^57^, Michelle Wantoch ^86^, Joanne Watts ^48^, Sheila Waugh ^66^, Sam Weeks^41^, Rebecca Williams^31^, Iona Willingham^56^, Emma L Wise ^25, 29, 31^, Victoria Wright ^54^, Sarah Wyllie ^70^, and Jamie Young ^3^.

**Software and analysis tools**

Amy Gaskin^33^, Will Rowe ^15^, and Igor Siveroni ^96^.

**Visualisation**

Robert Johnson ^96^.

**1** Wellcome Sanger Institute, **2** Public Health England, **3** University of Cambridge, **4** Health Data Research UK, Cambridge, **5** Public Health Agency, Northern Ireland, **6** Queen’s University Belfast **7** Public Health England Colindale, **8** Department of Medicine, University of Cambridge, **9** University of Oxford, **10** Departments of Infectious Diseases and Microbiology, Cambridge University Hospitals NHS Foundation Trust; Cambridge, UK, **11** Division of Virology, Department of Pathology, University of Cambridge, **12** The Francis Crick Institute, **13** Cambridge Institute for Therapeutic Immunology and Infectious Disease, Department of Medicine, **14** Public Health England, Clinical Microbiology and Public Health Laboratory, Cambridge, UK, **15** Institute of Microbiology and Infection, University of Birmingham, **16** University of Birmingham, **17** Queen Elizabeth Hospital, **18** Heartlands Hospital, **19** University of Edinburgh, **20** NHS Lothian, **21** MRC-University of Glasgow Centre for Virus Research, **22** Institute of Biodiversity, Animal Health & Comparative Medicine, University of Glasgow, **23** West of Scotland Specialist Virology Centre, **24** Dept Zoology, University of Oxford, **25** University of Surrey, **26** Wellcome Centre for Human Genetics, Nuffield Department of Medicine, University of Oxford, **27** Big Data Institute, Nuffield Department of Medicine, University of Oxford, **28** Oxford University Hospitals NHS Foundation Trust, **29** Basingstoke Hospital, **30** Centre for Genomic Pathogen Surveillance, University of Oxford, **31** Hampshire Hospitals NHS Foundation Trust, **32** University of Southampton, **33** Public Health Wales NHS Trust, **34** Cardiff University, **35** Betsi Cadwaladr University Health Board, **36** Cardiff and Vale University Health Board, **37** Swansea University, **38** University of Sheffield, **39** Sheffield Teaching Hospitals, **40** Great Ormond Street NHS Foundation Trust, **41** University College London, **42** Oswaldo Cruz Institute, Rio de Janeiro **43** North West London Pathology, **44** Imperial College Healthcare NHS Trust, **45** NIHR Health Protection Research Unit in HCAI and AMR, Imperial College London, **46** University of Liverpool, **47** Manchester University NHS Foundation Trust, **48** Liverpool Clinical Laboratories, **49** University of Exeter, **50** Royal Devon and Exeter NHS Foundation Trust, **51** Quadram Institute Bioscience, University of East Anglia, **52** Norfolk and Norwich University Hospital, **53** University of East Anglia, **54** Deep Seq, School of Life Sciences, Queens Medical Centre, University of Nottingham, **55** Virology, School of Life Sciences, Queens Medical Centre, University of Nottingham, **56** Clinical Microbiology Department, Queens Medical Centre, **57** PathLinks, Northern Lincolnshire & Goole NHS Foundation Trust, **58** Clinical Microbiology, University Hospitals of Leicester NHS Trust, **59** Viapath, **60** Hub for Biotechnology in the Built Environment, Northumbria University, **61** NU-OMICS Northumbria University, **62** Northumbria University, **63** South Tees Hospitals NHS Foundation Trust, **64** North Cumbria Integrated Care NHS Foundation Trust, **65** North Tees and Hartlepool NHS Foundation Trust, **66** Newcastle Hospitals NHS Foundation Trust, **67** County Durham and Darlington NHS Foundation Trust, **68** Centre for Enzyme Innovation, University of Portsmouth, **69** School of Biological Sciences, University of Portsmouth, **70** Portsmouth Hospitals NHS Trust, **71** University of Warwick, **72** University Hospitals Coventry and Warwickshire, **73** Warwick Medical School and Institute of Precision Diagnostics, Pathology, UHCW NHS Trust, **74** Genomics Innovation Unit, Guy’s and St. Thomas’ NHS Foundation Trust, **75** Centre for Clinical Infection & Diagnostics Research, St. Thomas’ Hospital and Kings College London, **76** Department of Infectious Diseases, King’s College London, **77** Guy’s and St. Thomas’ Hospitals NHS Foundation Trust, **78** Centre for Clinical Infection and Diagnostics Research, Department of Infectious Diseases, Guy’s and St Thomas’ NHS Foundation Trust, **79** Princess Alexandra Hospital Microbiology Dept., **80** Cambridge University Hospitals NHS Foundation Trust, **81** East Kent Hospitals University NHS Foundation Trust, **82** University of Kent, **83** Gloucestershire Hospitals NHS Foundation Trust, **84** Department of Microbiology, Kettering General Hospital, **85** National Infection Service, PHE and Leeds Teaching Hospitals Trust, **86** Cambridge Stem Cell Institute, University of Cambridge, **87** Public Health Scotland, 88 Belfast Health & Social Care Trust, **89** Health Services Laboratories, **90** Barking, Havering and Redbridge University Hospitals NHS Trust, **91** Royal Free NHS Trust, **92** Maidstone and Tunbridge Wells NHS Trust, **93** University of Brighton, **94** Kings College London, **95** PHE Heartlands, **96** Imperial College London, **97** Department of Infection Biology, London School of Hygiene and Tropical Medicine.

## Notes

### Competing Interest Statement

The authors have declared no competing interest.

## References

1. Gardy JL, Loman NJ. Towards a genomics-informed, real-time, global pathogen surveillance system. Nat Rev Genet. 2018;19: 9–20.

2. The COVID-19 Genomics UK (COG-UK) consortium. An integrated national scale SARS-CoV-2 genomic surveillance network. The Lancet Microbe. 2020;1: e99–e100.

3. Connor TR, Loman NJ, Thompson S, Smith A, Southgate J, Poplawski R, et al. CLIMB (the Cloud Infrastructure for Microbial Bioinformatics): an online resource for the medical microbiology community. Microb Genom. 2016;2: e000086.

4. Quick J, Loman NJ, Duraffour S, Simpson JT, Severi E, Cowley L, et al. Real-time, portable genome sequencing for Ebola surveillance. Nature. 2016;530: 228–232.

5. Wu F, Zhao S, Yu B, Chen Y-M, Wang W, Song Z-G, et al. A new coronavirus associated with human respiratory disease in China. Nature. 2020;579: 265–269.

6. Black A, MacCannell DR, Sibley TR, Bedford T. Ten recommendations for supporting open pathogen genomic analysis in public health. Nat Med. 2020;26: 832–841.

7. Griffiths EJ, Timme RE, Page AJ, Alikhan N-F, Fornika D, Maguire F, et al. The PHA4GE SARS-CoV-2 Contextual Data Specification for Open Genomic Epidemiology. other. 2020. doi: 10.20944/preprints202008.0220.v1

8. Django Software Foundation. Django. Available: https://djangoproject.com

9. Di Tommaso P, Chatzou M, Floden EW, Barja PP, Palumbo E, Notredame C. Nextflow enables reproducible computational workflows. Nat Biotechnol. 2017;35: 316–319.

10. Köster J, Rahmann S. Snakemake--a scalable bioinformatics workflow engine. Bioinformatics. 2012;28: 2520–2522.

11. Argimón S, Abudahab K, Goater RJE, Fedosejev A, Bhai J, Glasner C, et al. Microreact: visualizing and sharing data for genomic epidemiology and phylogeography. Microb Genom. 2016;2: e000093.

12. Shu Y, McCauley J. GISAID: Global initiative on sharing all influenza data - from vision to reality. Euro Surveill. 2017;22. doi: 10.2807/1560-7917.ES.2017.22.13.30494

